# Probabilistic migration events drive transient tissue residency of lymphocytes during homeostasis

**DOI:** 10.1101/2025.07.04.663225

**Authors:** Václav Gergelits, Oliver T. Burton, James Dooley, Carlos P. Roca, Orian Bricard, Arman Ghodsinia, Emanuela Pasciuto, Vikram Sunkara, Adrian Liston

## Abstract

Tissue-resident lymphocytes form a phenotypically and functionally distinct analog to the corresponding circulatory lymphocyte populations. Residential CD8 T cells, in particular, are identified as having prolonged residence in the tissues and key functions in recall responses at tissue-environmental interfaces, although the dwell time in individual tissues has yet to be resolved. Residential CD4 T cells, regulatory T cells, B cells, and NK cells have been demonstrated to share phenotypic properties with residential CD8 T cells, but the migratory kinetics are even more poorly defined. Here we used probabilistic modelling on a large parabiosis dataset, covering multiple time-points and tissues, to calculate migration kinetics and dwell times of multiple lymphocyte subsets across a diverse set of tissues. Markov chain modelling identified distinct cell type-specific and tissue-specific residency patterns. The liver and gut were prone to prolonged residency compared to other tissue types, and a hierarchy of residency was observed with CD8 T cells and NK cells demonstrating longer residency than CD4 conventional T cells and regulatory T cells, which in turn resided in tissues longer than B cells. With few exceptions, however, average residency was at least an order of magnitude shorter than the life-span of the mouse, indicating a more dynamic form of steady-state tissue residency than usually assumed. Together these data provide a comprehensive model of a pan-tissue shared program in lymphocyte tissue residence, as well as identifying cell type- and organ-specific modification of the migratory kinetics.

## Introduction

Beyond the lymphocyte populations present in the blood and lymphoid organs, a population of tissue-resident lymphocytes is found within most tissues. These tissue-resident lymphocytes often exhibit unique properties and functions, in addition to their altered anatomical residence. The best studied example of tissue-residency remains the tissue resident memory (TRM) CD8 T cell population, which have been demonstrated to play a critical role in recall responses against infections in the brain ^1^, skin ^2, 3^, vagina ^2, 4^, lung ^5, 6^, liver ^7^ and gut ^8^. Resident CD8 T cells may also have benefits in anti-tumor responses ^9^, while, conversely, accentuating graft-vs-host disease ^10^. TRM CD8 T cells have a distinct phenotypic and transcriptional profile ^11, 12^, with CD69 the best individual marker correlating with tissue residency ^13^. The transcriptional and epigenetic residency module is conserved beyond CD8 TRM, correlating with the markers upregulated in other tissue-resident lymphocytes ^14^. While less studied, the critical functions of these other resident lymphocyte populations are also increasingly being recognised, with tissue-resident conventional CD4 T cells (Tconv) contributing to protective responses against infections and cancer ^15, 16, 17, 18^ and tissue-resident regulatory T cells (Tregs) having diverse reparative and anti-inflammatory roles ^19, 20, 21, 22^. Innate lymphoid cells, such as NK cells, are also found across the tissues ^23^, and even tissue-resident B cells are now recognised and functionally-important ^24^.

With tissue-residency being the defining feature of the lineage, extensive efforts have gone into defining the cellular kinetics of the resident population. The primary tool for identifying the resident lymphocyte population within the tissue has been parabiosis, in which the circulatory systems of congenic mouse strains are surgically united, allowing a distinction to be made between populations that have entered the tissue prior or post the surgical fusion.

Parabiotic studies of tissue-resident lymphocytes have been performed under steady-state homeostatic conditions, or following challenges such as infection, transplantation or immunogen challenge ^3, 15, 23, 25, 26, 27^. In the vast majority of the parabiotic studies, only a single time was used, an approach which provides formal proof of extended residency of at least a fraction of the population present within the tissue, and which enables the phenotypic definition of the population with longer residency ^28^. This approach has been successfully used to deconstruct the molecular requirements for tissue residency of lymphocyte subsets ^13^. Single time point parabiotic analysis has the key limitation of treating the population as bifurcated, between the transient population replaced since surgery and the residual population often assumed to have indefinite residency. When parabionts from multiple timepoints after parabiosis have been analysed, they demonstrate a continual rather than bifurcated population structure ^3, 29, 30, 31^, although without further mathematical modelling to provide deeper understanding of the tissue residency lymphocyte dynamics. Conversely, a plethora of advanced mathematical models have been applied to the study of lymphocyte residency, including differential equations and random walks ^32, 33^. These modelling approaches have, however, been limited to data more suited for short-term cell tracing experiments, and typically focused on the dynamics of naïve lymphocytes within secondary lymphoid organs, rather than tackling the question of lymphocyte residency within the non-lymphoid tissues.

With the advantages of parabiosis experiments in tracing cell migration and phenotype change across the prolonged time-course required to assess residency in the non-lymphoid tissues, there have been calls to apply advanced mathematical modelling to parabiotic data to understand tissue lymphocyte kinetics ^34^. Here we performed probabilistic computational modelling on a large-scale parabiotic dataset, covering multiple lymphocyte lineages (CD8 T cells, Tconv, Tregs, B cells, NK cells) with multiple time-points and a pan-tissue perspective. The comprehensive nature of the dataset allowed us to develop a Markov chain model of the cellular migration, differentiation and survival kinetics of both each lymphocyte lineage, with estimates of the rate of cellular transition events across the different organs. This model identifies a highly dynamic process of tissue residency for lymphocytes under homeostatic conditions, with the dwell time of lymphocyte lineages within particular organs derived from the probabilistic replacement rates driving continual turnover.

## Results

### Develop a model for T cell dynamics across lymphoid and non-lymphoid tissues

In order to understand the population dynamics of T cell residency in the tissues at the equilibrium state, we performed in-depth mathematical modelling of a large multi-timepoint parabiotic experiment ^31^, consisting of CD45.1 mice parabiosed to CD45.2 mice, and assessed at 0, 1, 2, 4, 8 and 12 weeks post-surgery (**Figure 1A**). At each timepoint, the host/donor composition was assessed in the blood, lymphoid tissues (bone-marrow, spleen, lymph nodes, mesenteric lymph nodes and Peyer’s patches), non-lymphoid tissues (adrenals, brain, kidney, lung, liver, muscle, white adipose tissue, skin, pancreas, and female reproductive tract) and the gut (intraepithelial, lamina propria, and Peyer’s patches). High dimensional flow cytometry was previously used to identify Foxp3^+^ Tregs ^31^, and was here used to identify CD4 and CD8 conventional (Foxp3^-^) T cells, with the key phenotypic states being classified as naïve (CD44^-^CD62L^hi^), activated / antigen-experienced (CD44^+^CD62L^low^CD69^-^) and CD69^+^. To develop the framework for modelling T cell dynamic equilibrium, we used Markov chain modelling of kinetics, where the cell population within each individual tissue was modelled in an interconnected population set consisting of the tissue, the blood, and a combined pool of all other tissues, with the corresponding population sets of the parabiotic pairs being linked only through the blood, resulting in eighteen cellular states present in each parabiotic pair (**Figure 1B**).

**Figure 1.**
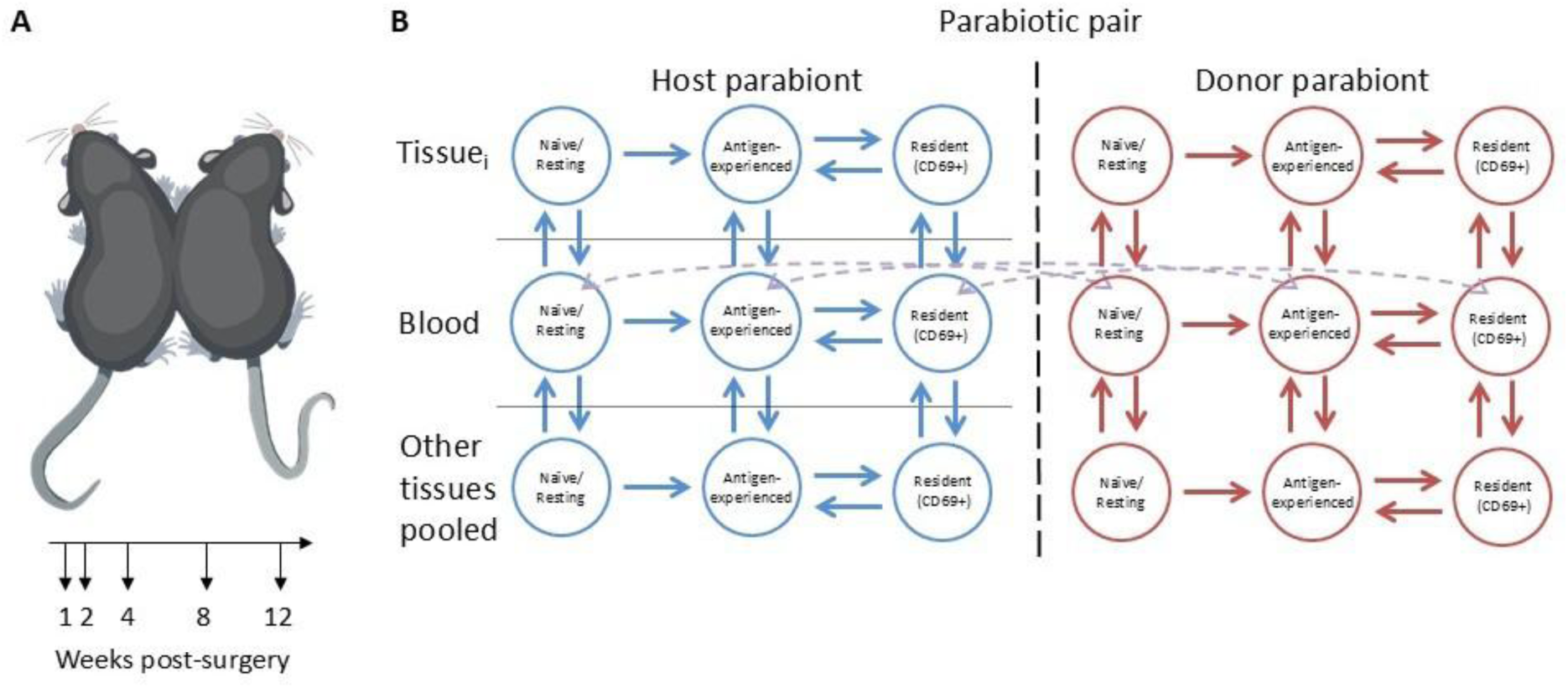
Markov chain modelling of T cell parabiosis data. **(A)** Pairs of parabiotic animals were sacrificed at weeks 1, 2, 4, 8, and 12 for tissue analysis. **(B)** The CD8 and CD4 T cells parabiosis data were modelled using closed Markov chains with 18 states representing naïve, activated and resident T cells in each tissue (once for the tissue of interest, once for the combined set of all other tissues) and naïve and activated T cells in the blood for both mice in the pair, with permissible transitions indicated. The models of this structure were estimated separately for each tissue in a closed system with blood.

Differentiation between the naïve and antigen-experienced states was considered irreversible, based on the standard biological model, while all other transitions were permitted. The parameters estimating different cellular processes of activation, migration and death were constrained by upper limits placed higher than plausible biology (Methods) to increase the speed of Markov Chain Monte Carlo (MCMC) sampling. As MCMC samples and best fit rates were orders-of-magnitude lower than the boundary limits, imposition of these limits did not alter the best fit models. Under these constraints, a Markov chain probabilistic model was built and estimated using Bayesian analysis, allowing estimation of the model transition rates and, consequently, residence times. Each Markov chain model was estimated by four Monte Carlo chains, with 5500 samples per chain including 500 samples of burn-in. For each transition rate, the mode posterior value is reported, together with the 80% highest density interval (Bayesian credibility). This approach allows a comparison of the transition rates between different cell types and different tissues, most likely to fit the empirical data observed.

### CD8 T cell migration dynamics change across tissue type

Using the model that fit the empirical data (**Figure 2A**, **Supplementary Figure 1**) with highest confidence, we first assessed the key transition rates for CD8 T cells across different tissues. The model predicted high entry rates (∼10^4^-10^5^ events / 1000 cells / day) for naïve (**Figure 2B**), activated (**Figure 2C**) and CD69^+^ (**Figure 2D**) CD8 T cells to enter lymphoid tissues (spleen, lymph nodes, mesenteric lymph node and bone-marrow). Entry into gut-associated tissues was more than 10-fold lower (∼10^2^-10^4^ events / 1000 cells / day), while non-lymphoid non-gut tissues ranged at the top end to similar to the gut (muscle, skin, pancreas) down to rates more than 1000-fold lower than lymphoid tissues (∼10^0-102^ events / 1000 cells / day). Other than the liver and Peyer’s Patches, naïve CD8 T cells had lower entry rates to non-lymphoid tissues than their antigen-experienced counterparts. Population flow diagrams demonstrate the interconnection of entry flows to differentiation, death and exit flows (**Figure 2E**). These present the stability of the population sizes within the different tissue compartments masking highly dynamic fluxes, which differ based on the cellular compartment. The lymphoid tissues included both highly dynamic migration fluxes of all cell states, plus the majority of naïve to activation transition events in the system. The non-lymphoid tissues, by contrast, had proportionally the highest entry of activated cells and conversion to CD69^+^ populations (**Figure 2E**). The best fit model also allowed the comparison of modelled dwell times for each subset across tissues (**Figure 2F**). Here a clear hierarchy was observed, with the longest dwell times recorded for CD69^+^ CD8 T cells in the gut-associated tissues, with a mean dwell time of 122 days in the IEL/LPL populations.

**Figure 2.**
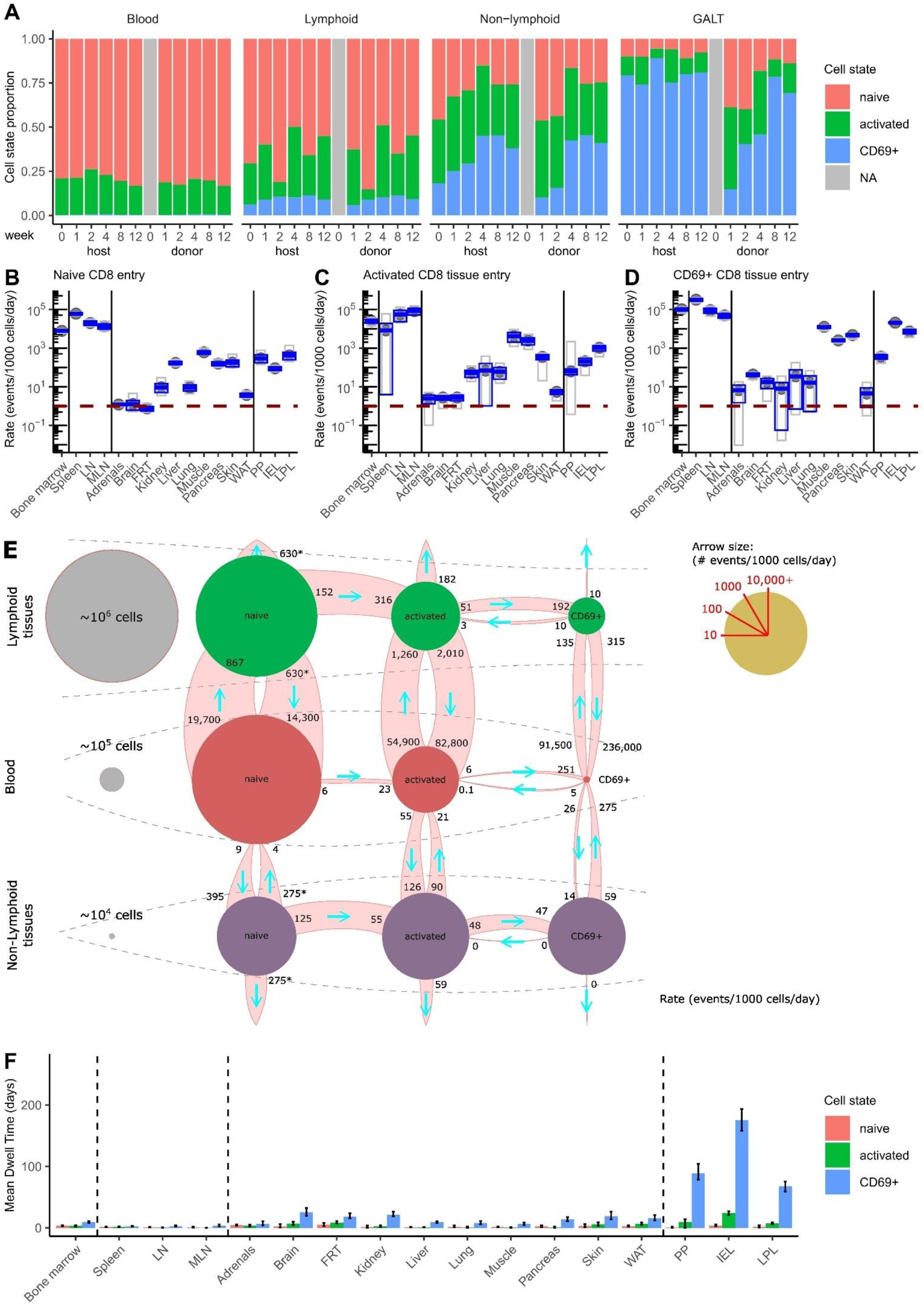
Differential kinetics for CD8 T cell migration in lymphoid vs non-lymphoid tissues. **(A)** Parabiotic data for CD8 T cell subsets, indicating the fraction of host- and donor-derived cells that are naïve, activated or CD69+. Data is displayed as blood, average of lymphoid tissues (bone marrow, spleen, LN, MLN), average of non-lymphoid (adrenals, brain, kidney, FRT, lung, liver, muscle, WAT, skin, pancreas) and average of gut-associated tissues (LPL, IEL, PP). **(B)** Markov chain modelling was used to estimate CD8 T cell flow/transition rates (measured by the number of events per 1000 cells per day) for lymphoid, non-lymphoid and gut tissue entry flow rate of naïve CD8 cells, **(C)** activated CD8 T cells, and **(D)** CD69+ CD8 T cells. **(E)** Model best fit CD8 T cell flow/transition rates (measured by the number of events per 1000 cells per day). For each tissue type (lymphoid, non-lymphoid, gut-associated), the median model result for each individual tissue, plus the median of the different tissues of that type, is displayed. Results are shown for naïve, activated and CD69+ cells, for tissue entry rates, tissue exit rates, apoptosis rates, differentiation (naïve to activated, activated to CD69+) and de-differentiation (CD69+ to activated). **(F)** Calculation of mean dwell time for CD8 T cells within each tissue, for the naïve, activated and CD69+ subsets.

Dwell times for CD69^+^ CD8 T cells were shorter in the non-lymphoid non-gut tissues (15 days), and shorter again for the lymphoid tissues (only 3 days). Naïve and activated CD8 T cells had shorter dwell times across the tissues, consistent with the status of CD69 as a residency marker.

### Conventional CD4 T cell dynamics across tissue source

Next, we analyzed the best-fit model for the CD4 Tconv cell parabiosis data (**Figure 3A**, **Supplementary Figure 2**). As with CD8 T cells, entry rates for CD4 Tconv were highest (10^3^-10^5^ events / 1000 cells / day) into lymphoid tissues, across naïve (**Figure 3B**), activated (**Figure 3C**) and CD69^+^ (**Figure 3D**). Gut entry rates were ∼100-fold lower, with a large range, from naïve cells at 10^1^-10^3^ events / 1000 cells / day to CD69^+^ cells at ∼10^4^ events / 1000 cells / day. Non-gut non-lymphoid tissues ranged from the highest entry tissues (liver, lung, muscle, pancreas) with entry rates similar to the gut tissues, to tissues such as the brain with rates around a hundred lower (∼1 event / 1000 cells / day). Among the non-lymphoid tissues, the liver demonstrated the best relative entry rate for naïve CD4 Tconv, with entry rates otherwise generally lower than the activated and CD69^+^ counterparts. The population flow diagrams demonstrate the interplay of entry, differentiation, death and exit flows, which indicate highly dynamic migration between blood and lymphoid tissues, with the majority of activation events occurring in the lymphoid tissues, and the non-lymphoid tissues largely populated by incoming activated and CD69^+^ cells (**Figure 3E**). For CD4 Tconv cells, the highest dwell times in each tissue were observed for the CD69^+^ cells, with the liver (>200 days) and bone-marrow (∼100 days) recording the highest dwell time for the CD69^+^ population. Other non-lymphoid tissues, including the gut, had mean dwell times ∼50 days, with lymphoid tissue dwell times substantially shorter (**Figure 3F**).

**Figure 3.**
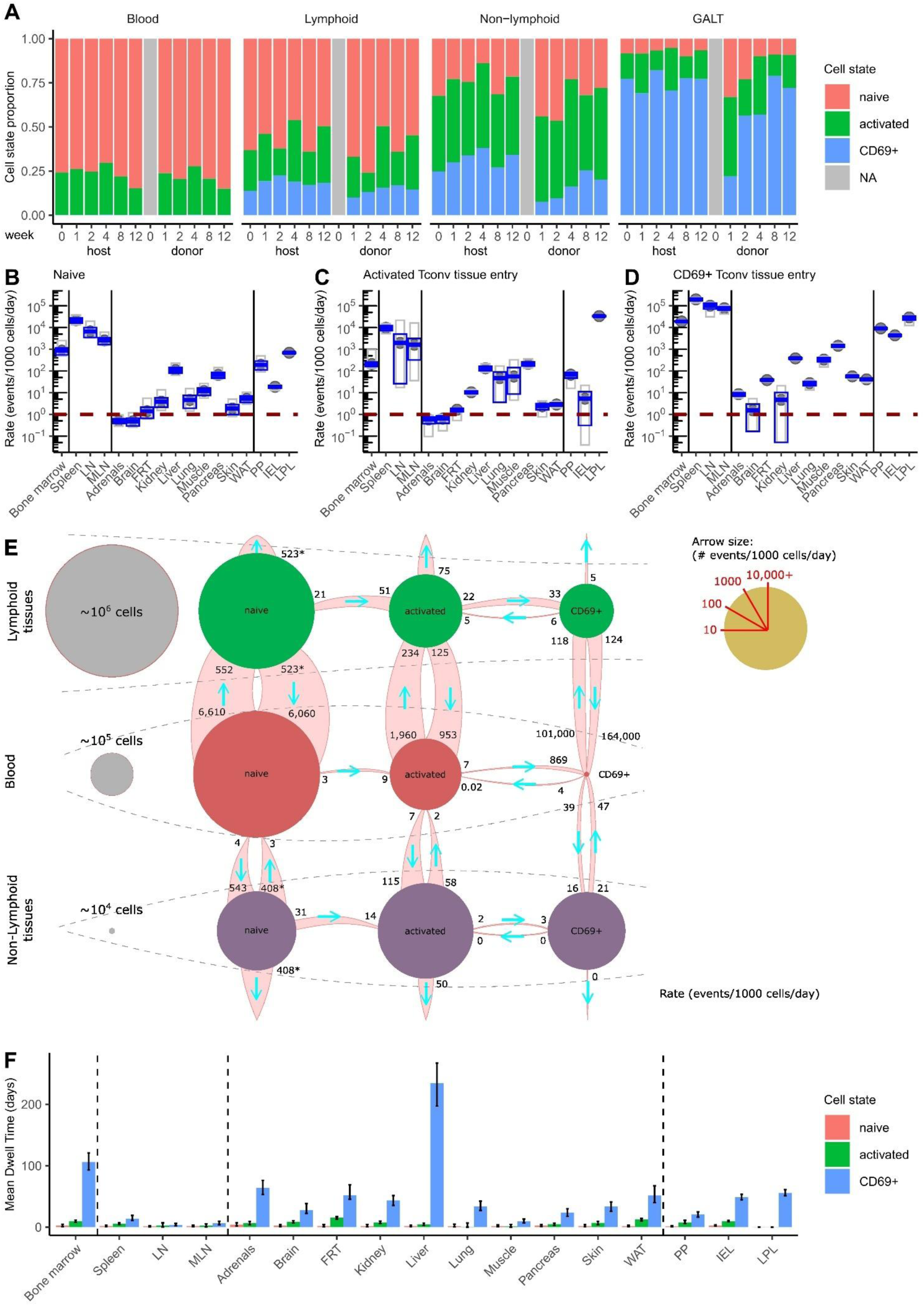
Differential kinetics for CD4 conventional T cell migration in lymphoid vs non-lymphoid tissues. **(A)** Parabiotic data for CD4 Tconv cell subsets, indicating the fraction of host- and donor-derived cells that are naïve, activated or CD69+. Data is displayed as blood, average of lymphoid tissues (bone marrow, spleen, LN, MLN), average of non-lymphoid (adrenals, brain, kidney, FRT, lung, liver, muscle, WAT, skin, pancreas) and average of gut-associated tissues (LPL, IEL, PP). **(B)** Markov chain modelling was used to estimate CD4 Tconv cell flow/transition rates (measured by the number of events per 1000 cells per day) for lymphoid, non-lymphoid and gut tissue entry flow rate of naïve CD4 Tconv cells, **(C)** activated CD4 Tconv cells, and **(D)** CD69+ CD4 Tconv cells. **(E)** Model best fit CD4 Tconv cell flow/transition rates (measured by the number of events per 1000 cells per day). For each tissue type (lymphoid, non-lymphoid, gut-associated), the median model result for each individual tissue, plus the median of the different tissues of that type, is displayed. Results are shown for naïve, activated and CD69+ cells, for tissue entry rates, tissue exit rates, apoptosis rates, differentiation (naïve to activated, activated to CD69+) and de-differentiation (CD69+ to activated). **(F)** Calculation of mean dwell time for CD4 Tconv cells within each tissue, for the naïve, activated and CD69+ subsets.

### Relative residency kinetics across T cell types

The inclusion of Treg parabiosis data, previously published ^31^ but reanalysed using the same method as CD8 T cells and CD4 Tconv cells (**Supplementary Figure 3**), allowed a comparison across T cell types from the same dataset. Similar trends were observed for entry and apoptosis rate predictions across the T cell lineages. For entry, a hierarchy of entry capacity was observed with lymphoid tissues the highest, followed by gut tissues, followed by non-lymphoid non-gut tissues (**Figure 4A**). CD69^+^ cells demonstrated the highest per-cell entry rates, across tissue types and T cell types, and while differences were observed in specific tissues, at the tissue-type level, no major differences were observed between Treg, CD8 T cells and CD4 Tconv cells (**Figure 4A**). It is important to note that while CD69^+^ cells had the highest per-cell entry rates, the relative scarcity of these cells in the peripheral blood results in numerically higher levels of migration from the naïve and activated T cell pools, across tissues and lineages (**Supplementary Figure 4**). Apoptosis rates could only be predicted for the activated and CD69^+^ subsets, as our Markov flows did not distinguish between tissue exit and apoptosis for naïve subsets, due to modelling as an open system replenished as new naïve thymic emigrants. When comparing the activated and CD69^+^ subsets, a large survival advantage was observed by CD69^+^ cells in the non-lymphoid tissues, across all T cell types, and in the bone-marrow for CD4 Tconv cells and the gut-associated tissues for Tregs and CD8 T cells (**Figure 4B**). The estimated rates of cell death for CD69^+^ T cells is such that the median cell is calculated to survive 132-345 days, with a substantial minority of cells surviving timeframes approximating the lifespan of the mouse, especially for CD4 Tconv cells (**Supplementary Figure 5**). This disparity between CD69^+^ T cell lifespan and tissue residency dwell time highlights the migratory nature of the population. An alternative way to visualize the population flows for each cell type and tissue source is through a destiny chart, indicating the fraction of cells leaving each tissue pool through apoptosis or migration, at different differentiation stages (**Figure 4C-E**). Here the changes in different population flows manifest in changes in typical cell destinies. For lymphoid tissues, the primary destination flows were outward flows of naïve T cells for CD8 T cells and CD4 Tconv cells, while outward migration of antigen-experienced and CD69^+^ cells were more important for Tregs (**Figure 4C-E**). In gut tissues, outward migration of CD69^+^ cells was the most common destiny across cell types. Non-gut non-lymphoid tissues, by contrast, varied, with CD8 T cells generally showing a higher importance of outward migration of activated or CD69^+^ cells (**Figure 4C**), while CD4 Tconv had a more balanced outflow via death and migration (**Figure 4D**) and Treg were somewhat intermediate (**Figure 4E**). Together, these results demonstrate the dynamic nature of T cell residency, and are suggestive of a pan-T cell set of residency rules, with relatively few exceptions (CD8 T cells in the gut, CD4 Tconv cells in the liver) to limited residency dwell times.

**Figure 4.**
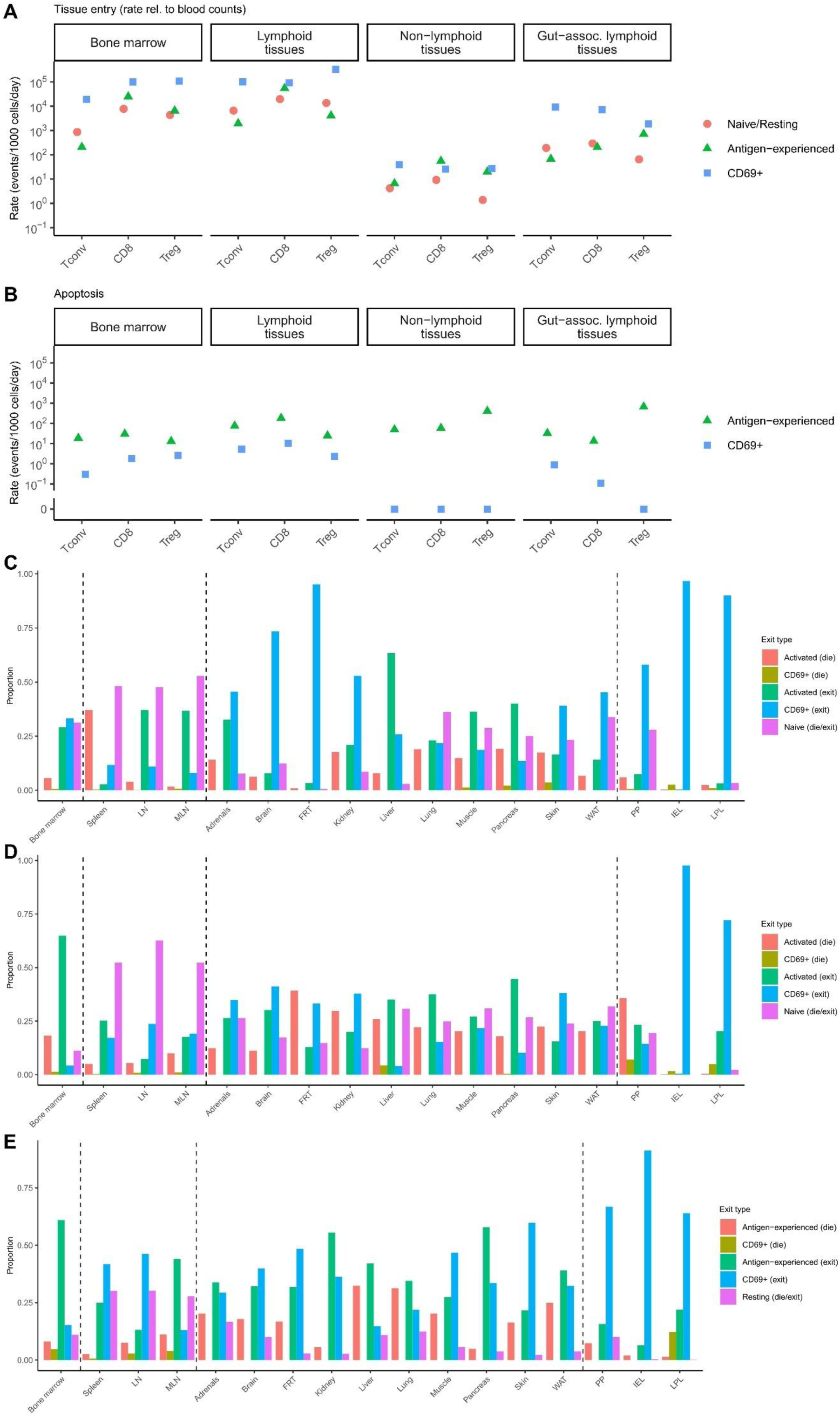
Tissue- and subset-specific variation in migration kinetics. **(A)** Best fit models from CD8 T cells, CD4 Tconv and CD4 Tregs were used to extract the tissue entry and **(B)** apoptosis rates for the naïve/resting, activated/antigen-experienced and CD69+ subsets. Rates were calculated for lymphoid (bone marrow, lymphoid tissues (spleen, LN, MLN), non-lymphoid (adrenals, brain, kidney, FRT, lung, liver, muscle, WAT, skin, pancreas) and gut-associated (LPL, IEL, PP) tissues. **(C)** Cell destiny, defined as the modelled pathway by which the cell terminated its residency in the tissue (exit or apoptosis, at each state of activation) was derived from the Markov chain model best fits for CD8 T cell, **(D)** CD4 Tconv, and **(E)** CD4 Treg cells.

### Highly divergent residency kinetics for B cells and NK cells

The flow cytometry markers for the experimental dataset ^31^ and the design of the Markov Chain modelling approach used here were both optimized for the analysis of T cells.

Nonetheless, markers were available to identify both B cells (CD19^+^) and NK cells (CD3^-^ NK1.1^+^) from the parabiotic pairs, enabling the MCMC analysis to be performed. We ran both cell types through the MCMC pipeline, which, despite the potentially sub-optimal use of naïve, activated and CD69^+^ cellular definitions, produced stable and robust Markov chains (**Supplementary Figure 6**, **Supplementary Figure 7**). B cell tissue residency was highly limited across tissues, with the CD69^+^ fraction in the IEL and LPL of the gut the only subsets to show residency of more than a few weeks, at ∼21 days (**Figure 5A**). NK cells, by contrast, had residency dwell times more similar to that observed in T cells, with typical dwell times for CD69^+^ cells of ∼50 days in gut and non-gut non-lymphoid tissues, extending out to ∼100 days in the female reproductive tract and the gut (**Figure 5B**). The lung was a notable exception to the prolonged tissue residency, consistent with prior studies ^23^, with residency shorter even than that observed in lymphoid tissues (**Figure 5B**).

**Figure 5.**
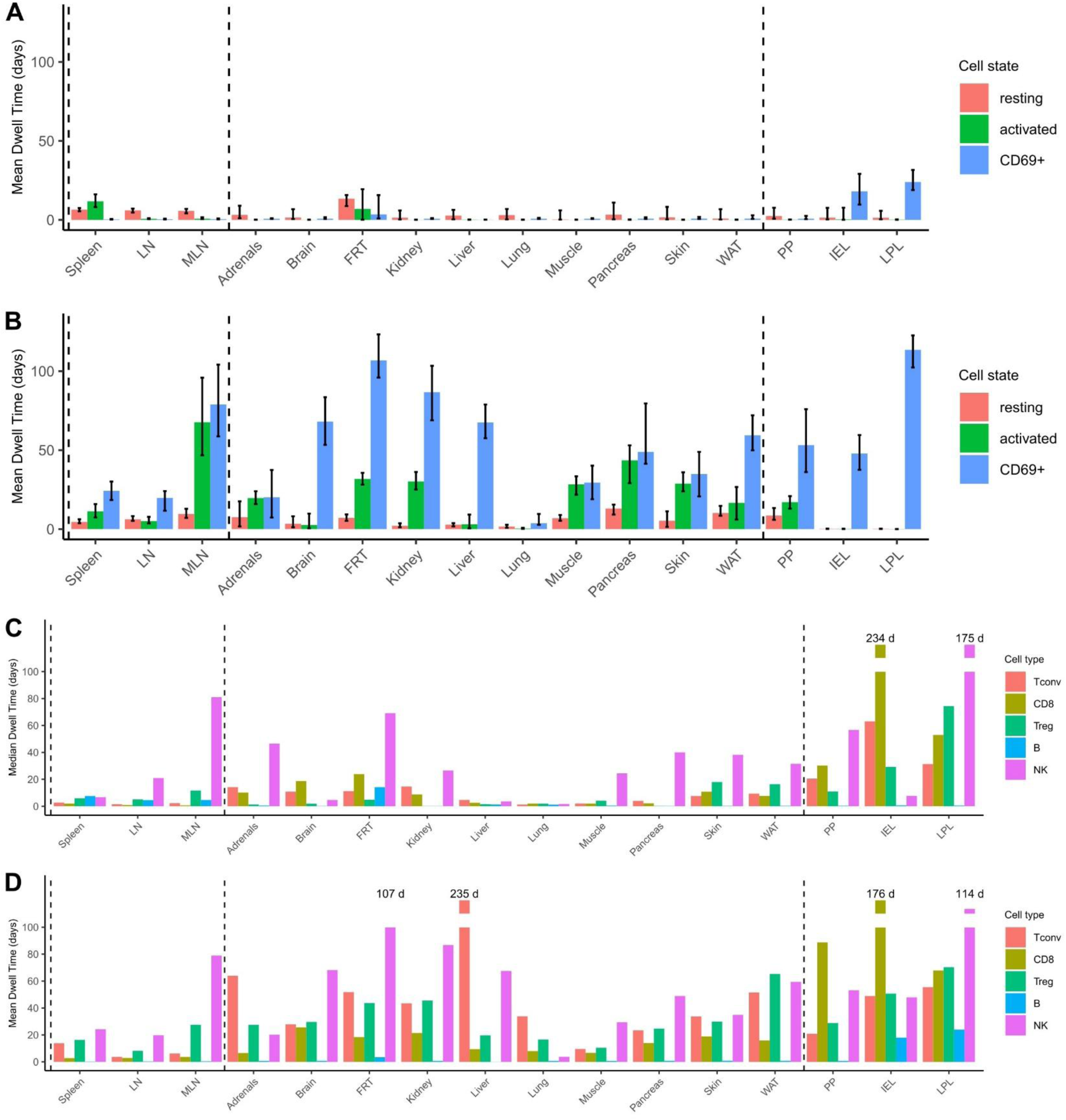
Broad utility of Markov chain modelling for lymphocyte subsets. **(A)** Markov chain modelling was used to estimate B cell and NK migration and transition rates (measured by the number of events per 1000 cells per day) for lymphoid tissues (spleen, LN, MLN), non-lymphoid tissues (adrenals, brain, kidney, FRT, lung, liver, muscle, WAT, skin, pancreas) and gut-associated tissues (LPL, IEL, PP). Best fit models were used to calculate mean dwell time for B cells and **(B)** NK cells, within each tissue, for the naïve, activated and CD69+ subsets. **(C)** Direct comparison of modelled lymphocyte dwell time across different tissues, for total population (median) and **(D)** CD69+ (mean) populations, for each of CD8, CD4 Tconv, CD4 Treg, B cells and NK cells.

Looking across the lymphocyte type measured here, we could measure duration of residency as either the mean residency of cells in the CD69^+^ state (the longest-lived assessed subset), or the median residency of the total population (taking into consideration the calculated differentiation dynamics and oscillation between CD69^+^ and activated states), with the latter function heavily dependent on the fraction of CD69^+^ cells within the tissue. When looking only at the total lymphocyte population, the pattern of lymphocyte residency that emerged was generally one of low dwell times in lymphoid tissues and for B cells generally, and modest dwell times of ∼1-3 weeks for T cells and NK cells across most non-lymphoid tissues (**Figure 5C**). Exceptions for longer dwell times were T cells in gut, in particular CD8 T cells in the IEL fraction of the gut, and NK cells in the LPL fraction of the gut and, to a lesser extent, in non-gut tissues such as the female reproductive tract (**Figure 5C**). The calculated dwell times for the CD69^+^ fraction, limited to the period for which the cell maintained CD69 expression, were generally higher across the tissues, with ∼2-6 weeks frequently observed for T cells, longer for NK cells and shorter for B cells. The largest shift in dwell times was for CD4 T cells in the liver, where the minor CD69+ fraction had very long residency, but was too small to substantially alter the total population dwell time (**Figure 5D**).

### Heterogeneity of residency emerges from probabilistic rules of lymphocyte migration, differentiation and death

Understanding the behaviour of the average cell (entry, apoptosis, dwell time, exit) sheds light on the dynamics of the system. However, a key feature of biological systems, and one which is replicated by Markov chains, is the probabilistic nature of events at the individual cellular level. The rates of these biological processes by individual cells dictates the average behaviour, however it does not represent the heterogeneity in cell fates that spontaneously emerges from probability-based systems. To illustrate the heterogeneity of fates within the populations, we generated *in silico* populations of “synthetic cells” based on the probabilistic rules extracted from the empirical data (**Figure 6**). Each cell independently underwent migration, differentiation and apoptosis events using the same probabilities, extracted from the relevant lymphocyte category. To visualize the heterogeneity within the tissue-resident populations, we plotted the residency fates of 1000 representative cells within each of the spleen, brain, liver, lung, white adipose tissue and gut IEL populations. Population dynamics for CD8 T cells demonstrated the diverse fates present across different organs (**Figure 6A**). In the spleen, for example, the combination of relatively low conversion to the CD69^+^ state and relatively high exit rates resulted in few cells being maintained more than 2 weeks. Liver and lung also have relatively few CD69^+^ cells, although the prolonged dwell time of those cells resulted in an L-shaped curve, with a small fraction of cells lasting 5-6 weeks in the tissues. Brain, adipose and IEL, by contrast, had larger fractions of cells entering the CD69^+^ state, resulting in a more linear decay in residency at the population level, dictated by the dwell time of the CD69^+^ population (**Figure 6A**). A similar distribution of tissue heterogeneity was observed in CD4 Tconv cells (**Figure 6B**) and Treg (**Figure 6C**), although it was notable that in the liver the prolonged longevity of the liver CD4 Tconv CD69^+^ population resulted in an extreme L-shaped curve, with the majority of naïve and activated Tconv cells rapidly replaced, while the CD69^+^ cells had only very slow turnover. The turnover of the B cell population, by contrast, was largely driven by the non-residential state of the bulk of the population, with activated B cells in the spleen turning over in a slower near-linear fashion, while non-lymphoid tissues were largely populated by highly transient naïve B cell infiltration, giving a short population-level half-life. Only the IEL population demonstrated the L-shaped curve, where the small population of CD69^+^ B cells lasted 5-6 weeks (**Figure 6D**). Each of the lymphocyte turnover patterns described above was also observed by NK cells, with the lung giving a rapid turnover of naïve and activated NK cells, similar to that observed in B cells, while delayed kinetics were observed in the spleen and adipose tissue, and the brain, liver and IEL populations followed L-shaped curves, with the CD69^+^ fraction maintained for weeks, more akin to the T cell model (**Figure 6E**). Indeed, for some lymphocyte populations, the most suitable time-scale involved hundreds of days (**Figure 6F**), with a small minority of CD4 Tconv cells and NK cells in the liver, and CD8 T cells, CD4 Tconv cells and NK cells in the gut IEL population, having dwell times that were in the range of the lifetime of the organism. It is important to note that this heterogeneity in residency times was not dependent on cell-specific factors, such as clonality, as it occurred in a synthetic population of cells with identical starting rules, and was simply an emergent property of probabilistic-based rules.

**Figure 6.**
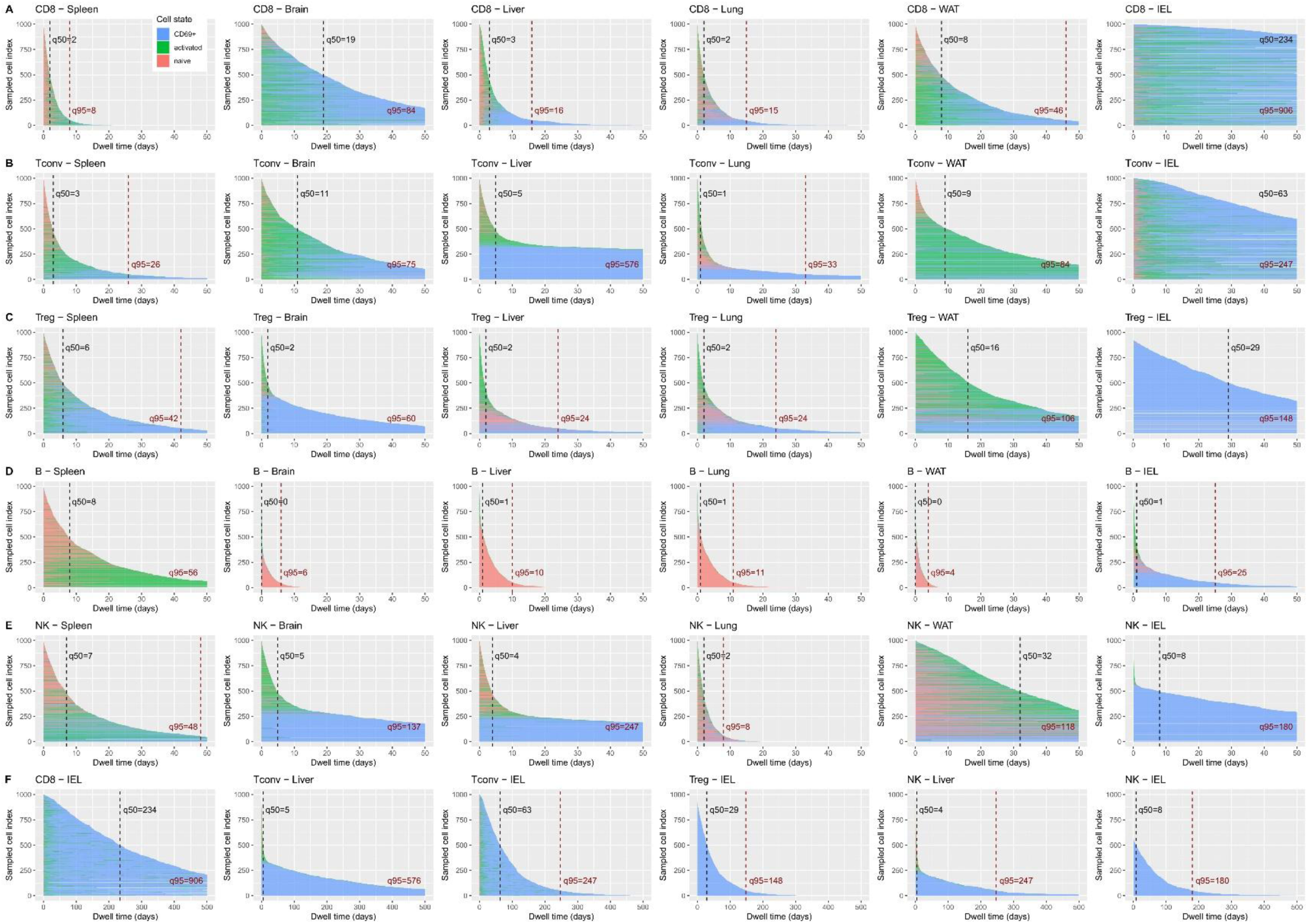
Heterogeneity in tissue residence emerges spontaneously from probabilistic rules. In order to model the implications of the cellular migration and state transition rules at the population level, “synthetic cells” were modelled *in silico*, as 1000 identical cells, each obeying the probabilistic rates identified in the best fit model, per lymphocyte lineage and per tissue. The duration and cell state of each cell is shown, ordered from the shortest modelled tissue residency through to the longest modelled tissue residency. Synthetic populations are shown for **(A)** CD8 **(B)** CD4 Tconv and **(C)** CD4 Treg cells **(D)** B cells and **(E)** NK cells, with **(F)** a zoom-out view on the lymphocyte-tissue combination with the longest total dwell times.

## Materials and Methods

### Source data generation

A series of parabiotic pairs were generated to study the migratory kinetics of tissue-resident regulatory T cells ^31^. In brief, pairs of 7-10 week-old *Foxp3^Thy^*^1.1^ female mice, C56BL/6 (CD45.2) and C57BL/6.SJL-Ptprca/BoyJ mice (CD45.1), were co-housed for 14-21 days prior to surgery. Pairs of mice were anesthetized with inhaled isoflurane, 3.5% v/v induction, 2.5-3.0% v/v maintenance. Carprofen and buprenorphine were delivered intraperitoneally at a dose of 10 mg/kg and 0.1 mg/kg prior to surgery. Mice were shaved and disinfected prior to longitudinal skin incisions, from 0.5cm above the elbow to 0.5cm below the knee joint. Detached skin was sutured together to generate a parabiotic pair. Pairs were euthanized at 1, 2, 4, 8, 12 weeks post-surgery, and assessed by flow cytometry of tissue preparations on a BD FACSymphony for CD45, CD45.1, CD4, CD8α, CD3, CD19, NK1.1, Foxp3, eBioscience™ Fixable Viability Dye eFluor™ 780, CD103, CD62L, CD25, Neuropilin, ST2, PD-1, CTLA4, KLRG1, Helios, CD69, ICOS, CD44, T-bet and Ki67. Data was compensated using AutoSpill ^35^. The raw flow cytometry data and analyzed data used as input for the parabiosis modelling, is deposited on Mendeley Data (doi: 10.17632/b4y3w9nbw2.1). Input data used for the parabiosis modelling, indicating the proportion of naïve (CD44^-^CD62L^hi^), antigen-experienced (CD44^+^CD62L^low^) and resident (CD69^+^) cells within the CD4^+^CD3^+^Foxp3^-^ and CD8^+^CD3^+^ populations, are available as **Supplementary Resource 1**.

### Data structure

The parabiotic data were collected at six different time points (0, 1, 2, 4, 8, 12 weeks) in each member of the parabiotic pair. Lymphoid tissues were considered to comprise pooled (non-mesenteric) lymph nodes (LN), mesenteric lymph nodes (MLN), spleen, and bone marrow (BM). Non-lymphoid tissues were comprised of adrenals, brain, kidney, liver, lung, muscle, pancreas, skin, white adipose tissue (WAT), and female reproductive tract (FRT). Gut-associated tissues were Payer’s patches (PP), intestinal intraepithelial lymphocytes (IEL) and lamina propria lymphocytes (LPL) . Blood was used as the reference point for each tissue.

For each time point and tissue, proportions of lymphocytes having specific cell surface marker combinations were estimated to define mouse of cell origin (host or donor, using CD45.1 or CD45.2 mismatching) and cell state (naïve, activated, resident). These proportions were estimated separately for conventional CD4 T cells, CD8 T cells, Tregs, NK cells and B cells.

### Probabilistic modelling

Parabiosis data was modelled using continuous-time Markov chain describing temporal evolution of average cell populations, while aiming to capture the actual mechanism governing lymphocyte behavior in mouse blood and tissues. The Bayesian approach was used to flexibly fit the complex structure of the Markov chain model and allowed for the direct generation of predictions about flow/transition rates of lymphocyte kinetics using their posterior distributions and dwell times in the model states. These predictions were before fitting restricted by conservative boundaries on the biologically plausible rates for each transition. The average cell populations were modeled with Dirichlet distributions and prior distributions were uninformative uniform from a wide range. To take advantage of available computational power we used efficient Markov chain Monte Carlo (MCMC) algorithm, specifically No-U-Turn Sampler (NUTS) variant of Hamiltonian Monte Carlo (HMC).

The dynamics of each lymphocyte subpopulation was modelled using independent continuous-time Markov chain models postulating that 1) individual lymphocytes behave independently, and 2) circuits between individual tested tissues, the blood and a compartment representing the rest of the body form independent and closed systems within the parabiotic pair of mice (**Figure 1B**). This latter assumption treats “blood” as the extra-tissue body composition, e.g. exit of lymphocytes via lymphatic drainage prior to reentering circulation would count as a tissue-to-blood transition. This gives (3 x 17) and (2 x 16) independently estimated models. The Markov chains were used to model the changes of lymphocyte states (naïve, activated, resident), and localization (blood, tissue, or combined pool of all other tissues) over time, with rates to change the state being estimated. The system was modelled to be in steady state, with the total number of lymphocytes dying in the system estimated to be equivalent to new naïve lymphocytes arising from the primary lymphoid organs, which was modelled as a circular system where dying lymphocytes transitioned to naïve lymphocytes in blood.

Conceptually, in the Bayesian modelling framework using the Bayes’ theorem: P(Q, **θ** | data) = P(data | Q, **θ**) × P(Q, **θ**) / P(data)

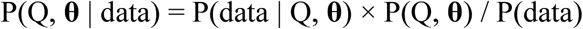

we were estimating the posterior distribution P(Q, **θ** | data ) of the set of flow/transition rate parameters in the intensity matrix Q characterizing the continuous-time Markov chain (**Figure 1B**) and nuisance vector **θ** described below. The likelihood P(data | Q, **θ**) at given parameters Q and **θ** correspond to observed data modelled using 18-dimensional Dirichlet distribution. The prior distribution P(Q, **θ**) was selected as uninformative.

For each Markov chain model, the data were modelled as probabilistic 18-dimensional simplex (2 mice x 9 subpopulations) with Dirichlet distribution evolving in time with mean values being averages of subpopulation proportions. We made the following restrictions to permissible transitions:

- Naïve lymphocytes can be activated but cannot return to the naïve state.
- Antigen-experienced lymphocytes can become resident, in a process the model permits to be reversible.
- The process of cell death cannot run concurrent to tissue migration. This technical boundary was not reached in any models.

All transition rates qij together formed an intensity (differential generator) matrix Q with 21 unknown free parameters (9 rates per 1 tissue + 9 rates per pool of other tissues + 3 host-donor rates). Furthermore, one free parameter to model total kinetics of the system was used while remaining non-zero parameters in matrix Q were calculated to fulfill the general probability requirements for intensity matrix. In addition, we used 15 free nuisance parameters (denoted as nuisance vector **θ** above) to capture the differential variability of observations in time and tissues (time points 1, 2, 4, 8, 12 weeks x 3 model parts).

The prior distributions for the free parameters were chosen as uninformative, uniform distribution between 0 and theoretically maximal rate in order not to provide any extra assumption that could bias the resulting posterior distribution. The maximal flow/transition rates were assumed in line with the assumptions above, specifically:

1) In tissues, the prior distribution of the free parameters representing cell-state transition rates of lymphocyte gain or loss of residency was assumed as uniform, U(0; 10,000) / 1000 cells / day. Furthermore, the lymphocyte activation rates were assumed to be bound by 10,000,000 events / 1000 cells / day.2) The prior distribution of activated and residential lymphocyte death rates was assumed as uniform, U(0; 5000) events / 1000 cells / day and U(0; 1000) events / 1000 cells / day, respectively.

3) The prior distribution of lymphocyte flow rates from tissue to blood was assumed as uniform, U(0; 10,000) events / 1000 cells / day for naïve lymphocytes, U(0; 10,000) events / 1000 cells / day for antigen-experienced lymphocytes and U(0; 100 / CD69_p) events / 1000 cells / day for residential cells, where CD69_p = max(limit CD69^+^ proportion in tissue, limit CD69^+^ proportion in other tissues combined).

4) The parameter of total outflux of naïve cell state in blood to naïve cells in combination of all tissues was constrained by initial boundary settings (500,000 for CD4 and 200,000 for CD8 T cells, 15,000 for NK cells, 4,000,000 for B cells) multiplied by a free parameter sampled from uniform distribution U(1, 30) to provide model initial whole-body dynamics estimate and to allow for subtle differences between models, each of which is focused on one tissue.

5) In blood, the prior distribution of all-state transition was assumed to be bound by 100 events / 1000 cells / day and assumed as uniform, U(0; 100) events / 1000 cells / day for loss of residency.

Generally, the upper boundary served as a necessary starting point to increase the speed of algorithms while the actual sampled values were orders of magnitude lower. This demonstrates that the results are not sensitive to the choice of prior distributions.

The exact form of model is available as a commented code in Stan language deposited at GitHub.

Each of the Markov chain model was estimated by 4 Monte Carlo chains with 5,500 samples (including 500 samples of burn-in) per chain. For each of the flow/transition rates the mode posterior value was reported together with 80% highest density interval (HDI), i.e. Bayesian, credibility interval spanning the most credible 80% of the estimated posterior distribution.

The Markov chain modelling performed by Markov Chain Monte Carlo estimation of Bayesian model posterior distribution was performed using the Stan probabilistic language. Namely, we run Stan using R language interface RStan in the form of R package rstan 2.32. Furthermore, we for other simulations and scripting we used R 4.5.1 and standard R packages, in particular expm, modeest, and tidyverse. The formatted input data is available as **Supplementary Resource 1**, the full code for analysis is provided at the GitHub repository https://github.com/gergelits/parabiosis.

## Discussion

In this study we have modelled the kinetics of lymphocyte residency in the tissues through the use of a probabilistic model. The approach fully recapitulates the complex population-level turnover that is experimentally observed, through the application of a limited set of cell behaviour traits (migration, activation, death) at the individual cell level. The multi-tissue and multi-lineage approach used for this modelling revealed heterogeneity in the cellular kinetics both between lineages (with CD8 T cells and NK cells having longer average residencies than CD4 Tconv and Treg cells, which in turn had longer residency than B cells) and across tissues (with the gut and liver showing the greatest duration of residency, for several lineages). Regardless of the combination of lineage and tissue, in each case the experimental data could be accurately modelled with probabilistic rates, where events such as emigration and death were continual processes with a linear cumulative probability of occurring over time. We did not require any subsets to operate by a “cellular clock” mechanism, whereby a fixed or minimum residency time would be applied, nor was there any residual signal that required a subset of indefinite duration to complete the model. These conclusions are in line even with classical experiments showing the maintenance of CD8 TRM against viral infections, which show a slow waning of residential cell frequency over time ^3^. The experimental data available is thus best explained by cells undergoing independent probabilistic decisions regarding migration, activation and death, with the rates of each decision altered by the local tissue environment.

The heterogeneity in tissue residency observed is intriguing. For the sake of interpretability, we have grouped individual tissues into types (lymphoid, gut-associated, non-gut non-lymphoid) based on anatomical location and our prior analysis of similarities between cell states ^31^. In general the gut-associated tissues had longer dwell-times than the non-gut tissues, and, with the exception of CD69+ CD4 Tconv cells in the liver, contained the longest dwell time of each lymphocyte lineage analysed. It is tempting to speculate, for T cells and B cells at least, that the uniquely high continual exposure to microbial and dietary antigens in the gut is responsible for this extended dwell time. The antigenic exposure is likely to reinforce the selection and survival of T cells and B cells bearing specific antigenic receptors against these antigens ^36, 37^. Alternatively, the longer residency observed here in the steady-state of the gut may reflect the normal physiological state of tissues, which, outside of SPF environments, bear a complex history of infection and inflammation that drive the formation of tertiary lymphoid structures that may alter lymphocyte migration ^38^. Likewise, multiple explanations are plausible for the different behaviour observed between lymphocyte lineages. The effects could be lineage-intrinsic, based on the developmental trajectories altering the cellular response to tissue microenvironments ^39^, or it could be based on different availability and maintenance of antigen exposure and presentation within the tissue post-infection ^40^. Further molecular insights into the nature of residency, comparing different lineages and tissues, will be needed to resolve these questions.

Many of the biological insights discussed above were not apparent from the primary data, which only provides information on whether host-donor equilibrium has been reached.

Rather, these insights required the application of mathematical modelling. In particular, Markov chain modelling was highly suited to the study of cellular kinetics, as it can infer steady-state population dynamics by fitting the properties of individual cells to the available time-point data. Increasing the complexity of migration based models to enable both cell-state transitions within tissues and migration between tissues, has been proposed to provide superior fits ^41^. Here, together with the multi-timepoint data series, it allowed more precise dwell time estimates in comparison to marginal models with only one source of cell influx considered. Our best fit multi-tissue models predicted cellular kinetic patterns that were identified through alternative methodologies (such as differential longevity in lymphoid tissues for naïve CD8 and CD4 Tconv cells ^42^ or the extended residency of CD69^+^ T cells ^43^), supporting the robustness and reliability of the approach. Other cellular kinetic features identified required the computational approach for derivation. These include the lower death rate of CD69^+^ T cells in the tissues, the extended lifespan of this subset, and the substantial recirculation of lymphocytes from the tissues back to the blood, which has only previously been inferred based on phenotype and clonal sharing ^43, 44^. The modelling approach also enables measurement of aspects of migration, such as exit rates, which have not been measured directly at the individual tissue level, and have only be estimated with difficulty at the organism level, such as through analysis of thoracic duct lymphocytes ^45^. Finally, the use of Markov chain modelling enables the generation of “synthetic cells” for in silico modelling. Such synthetic populations enable us to determine which heterogenous features require biological heterogeneity to explain (such as TCR or epigenetic clonality), and which, like variation in dwell time, can be explained purely through probabilistic processes, without invoking biological variation.

There are several important limitations to the current study. While we have developed a new computational framework for the analysis of parabiotic data, here we have only applied it to a dataset generated in the homeostatic state. Thus the conclusions reached about the cellular dynamics and transience of lymphocyte dwell-time in the tissues cannot be applied to lymphocyte behaviour in the post-infectious or inflamed tissue. There are limited reference points on whether the tissue-residency is shortened or extended following inflammation.

Virus-specific CD8 T cells, following influenza or vaccina infection, remained out-of-equilibrium in the parabiotic infected lung, at time points post-infection beyond the homeostatic normalization point identified here ^3, 27^, suggesting an extended dwell time. Likewise with CD8 and CD4 Tconv in the vagina following HSV infection ^15^. Even B cells, with the shortest tissue residency measured here in steady-state, can extend residency following infection, measuring influenza-reactive B cells in the parabiotic infected lung ^46^. Conversely, ILC2 cells showed enhanced turnover three weeks following helminth infection ^23^, as did tissue Tregs after inflammation of the omentum ^47^ and heart ^48^. Changes in lymphocyte tissue residency rules following infection are therefore not only expected, but are also likely to differ dramatically between lineages, and potentially across the tissue source and inflammatory stimulus. While the extended residency of post-infectious CD4, CD8 and B cell clones may be due to lymphocyte-intrinsic changes imparted during infection ^3, 27, 46^, it is also possible that local changes in the tissue efficiently capture clones in a tissue-led manner^15^.

Another set of limitations of our study are the simplifications made to enable effective modelling. First, we simplified the diversity of lymphocyte states to the naïve/resting, activated/antigen-experienced and CD69^+^/residential state. Additional relevant cell states may exist, with altered rules of cell dynamics. Furthermore, CD69 is unlikely to be a perfect marker for the population with enhanced residency, although deep profiling of the retained CD8 T cell parabiotic fraction suggests CD69 is the best single marker currently known ^13^, and phenotypic data supports the analogous use of this marker for Tregs ^31^, NK cells ^49^ and B cells ^50^. To overcome this potential limitation, we have provided here the original high-dimensional parabiosis data set, enabling independent assessment of subsets based on alternative candidate markers (see Data Availability). Regardless, any missing cell states must be of minor size or must have only minimal divergence from the modelled states, as the best-fit models generated here fit the empirical data well at the population level. Second, we simplified the anatomical connections to limit tissue-to-tissue transitions only via the blood. In many cases this simplification may accurately model the majority of transitions, however more interconnected models could be built up, for example allowing direct LPL and IEL interchange, and using the draining lymph node as a intermediate exchange point with the blood. While the current dataset does not have sufficient power to enable these more complex models to be assessed, the bioinformatic pipeline generated here is made available (see Code Availability) for application to analyse new parabiotic datasets generated to answer more specific questions on lymphocyte migration.

Finally, we need to consider the human context. Despite the difficulty of studying tissue-resident lymphocytes in humans, there has been a recent boom in studying this critical population ^51^. Tissue-resident lymphocytes are found across the tissues, with both shared and tissue-specific adaptations apparent, with heavy conservation towards the features identified in mouse tissue-resident lymphocytes ^14, 52, 53, 54, 55, 56, 57, 58^. Understanding which aspects of the cellular dynamics of tissue-residency are conserved across species and which will differ between mouse and human is critical for the therapeutic exploitation of these populations.

Only very limited data is available to feed into a kinetic model of lymphocyte residency in human tissues, for example the analysis of explanted transplantations. A seminal study of explanted liver transplants identified donor-derived tissue-resident CD8 T cells in the liver, even 10 years post-transplantation ^59^. Coupled with the low detection of donor-derived CD8 T cells in the hepatic vein and general circulation, this has been interpreted to reflect a degree of extreme longevity that is not present in the mouse homeostatic state (although the liver was among the organs with the longest dwell times). While this may reflect species differences, such discrepancy could also be explained by differences between the homeostatic state and the transplanted state, through the transplant-specific context of appropriate HLA being restricted only to the donated organ, or through circular migration providing a decoupling of detected residency from continual residency. Using steady-state from deceased donors and retrospective radiocarbon analysis of ^14^C from atmospheric nuclear weapons testing, tissue-resident CD8 T cells have extended lifetimes compared to the circulating population ^60^. This would be consistent both with the model of indefinite residency observed in human liver explant system ^59^, or with the results modelled here, where CD69^+^ CD8 T cells have high longevity and recirculation, rather than indefinite residency. Beyond CD8 T cells, human tissue-resident NK cells have been assessed through the analysis of menstrual blood in patients with either bone-marrow or uterine transplants ^61^. Here the data was consistent with rapid turnover of the tissue-resident NK population, in line with the murine modelling data. Further innovative studies will be required to determine the diversity of residency kinetics of lymphocyte lineages across different tissues in humans, in health and following infection or inflammation.

## Data availability

Source data for parabiosis flow cytometry time points, for conventional CD4 and CD8 T cells are provided as **Supplementary Resource 1**. The raw flow cytometry data is deposited on Mendeley Data (doi: 10.17632/b4y3w9nbw2.1).

## Code availability

Source code for parabiosis Markov chain modelling is available at the github repository https://github.com/gergelits/parabiosis.

## Supporting information

Supplementary resource 1

## Acknowledgements

This work was supported by the ERC Consolidator Grant TissueTreg (to A.L.) and the Wellcome Trust Investigator Award 222442/Z/21/Z (to A.L.). The authors declare no conflict of interests.

