## Supplementary material for "Probabilistic migration events drive transient tissue residency of lymphocytes during homeostasis"

**Supplementary Resource 1.** Source data for parabiosis flow cytometry time points, for both conventional CD4 and CD8 T cells, given as gated flow cytometry data.

**Supplementary Figure 1. Markov chain models for tissue CD8 cellular kinetics.** CD45.1 mice were parabiosed to CD45.2 mice. Pairs of parabiotic animals were sacrificed at weeks 1, 2, 4, 8, and 12 for tissue analysis by flow cytometry (n=11,12,18,16,14). Markov chains were built to model the changes in cell state and tissue exchange, with each tissue built using a model containing the tissue, blood, and combined other tissues. Displayed are the original data points superimposed on the model predictions for naïve CD8 (left), activated CD8 (middle) and $CD69^{+}$ CD8 (right) from **A.** bone-marrow, **B.** spleen, **C.** LN, **D.** mLN, **E.** adrenals, **F.** brain, **G.** FRT, **H.** kidney, **I.** liver, **J.** lung, **K.** muscle, **L.** pancreas, **M.** skin, **N.** WAT, **O.** PP, **P.** IEL, **Q.** LPL, and **R.** blood. The blood model results displayed is from the spleen, blood and other tissues model, while all other tissues are displayed from their own model.

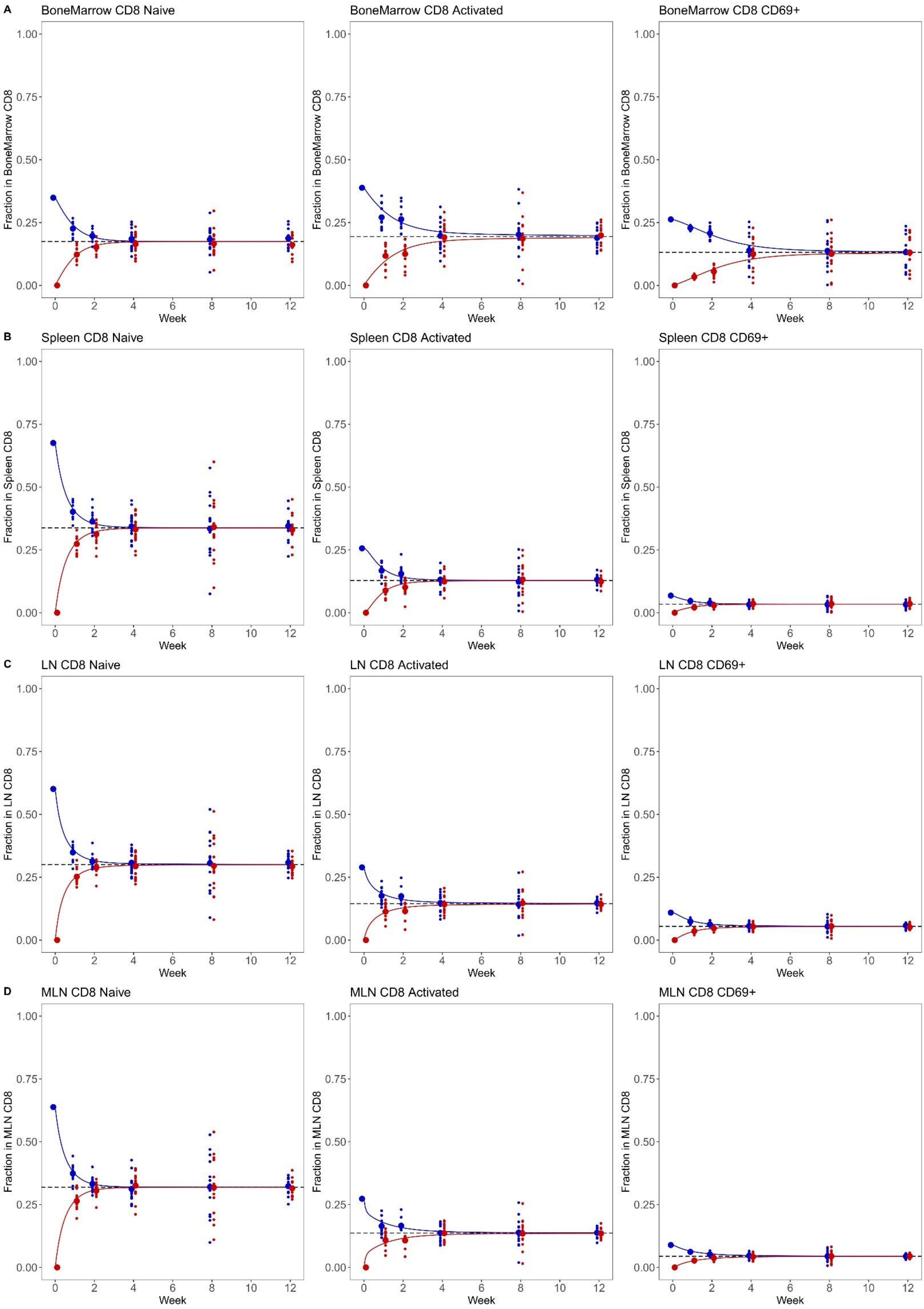

A
BoneMarrow CD8 Naive
BoneMarrow CD8 Activated
BoneMarrow CD8 CD69+
Fraction in BoneMarrow CD8
B
Spleen CD8 Naive
Spleen CD8 Activated
Spleen CD8 CD69+
Fraction in Spleen CD8
C
LN CD8 Naive
LN CD8 Activated
LN CD8 CD69+
Fraction in LN CD8
D
MLN CD8 Naive
MLN CD8 Activated
MLN CD8 CD69+
Fraction in MLN CD8
Week
0.00
0.25
0.50
0.75
1.00
0
2
4
6
8
10
12

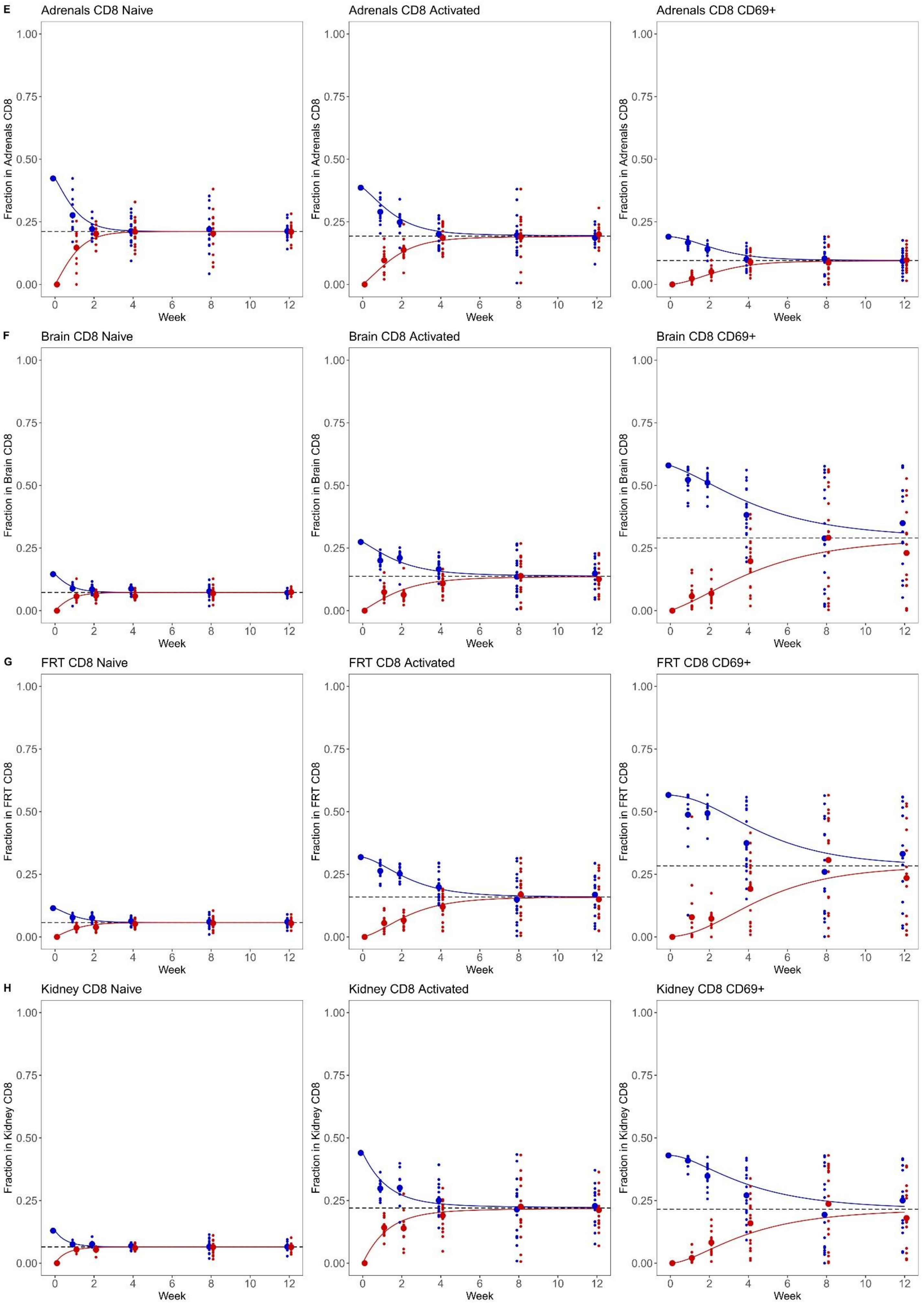

E
Adrenals CD8 Naive
Adrenals CD8 Activated
Adrenals CD8 CD69+
Fraction in Adrenals CD8
F
Brain CD8 Naive
Brain CD8 Activated
Brain CD8 CD69+
Fraction in Brain CD8
G
FRT CD8 Naive
FRT CD8 Activated
FRT CD8 CD69+
Fraction in FRT CD8
H
Kidney CD8 Naive
Kidney CD8 Activated
Kidney CD8 CD69+
Fraction in Kidney CD8
Week
0.00
0.25
0.50
0.75
1.00
0
2
4
6
8
10
12

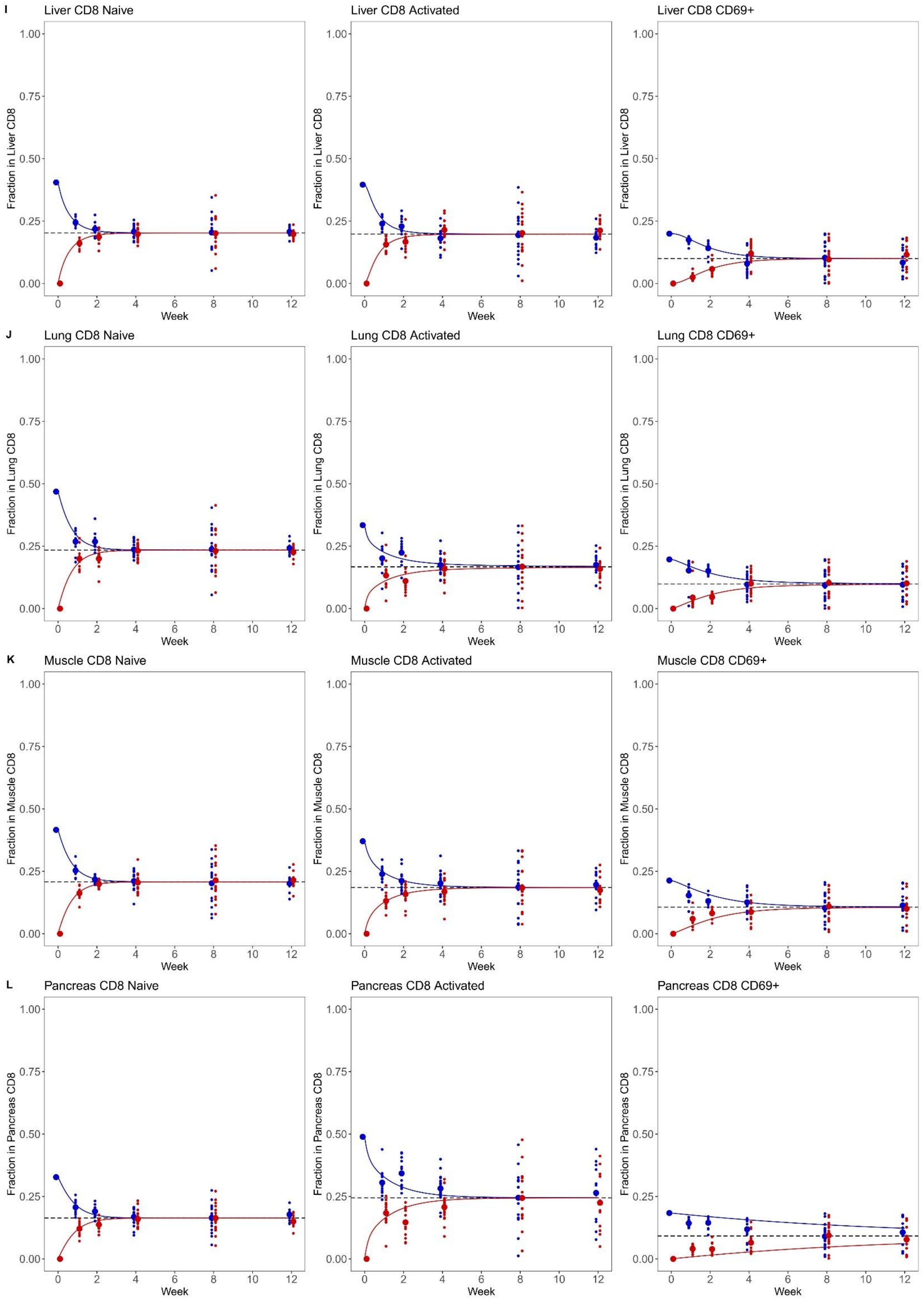

I
Liver CD8 Naive
Liver CD8 Activated
Liver CD8 CD69+
Fraction in Liver CD8
J
Lung CD8 Naive
Lung CD8 Activated
Lung CD8 CD69+
Fraction in Lung CD8
K
Muscle CD8 Naive
Muscle CD8 Activated
Muscle CD8 CD69+
Fraction in Muscle CD8
L
Pancreas CD8 Naive
Pancreas CD8 Activated
Pancreas CD8 CD69+
Fraction in Pancreas CD8
0.00
0.25
0.50
0.75
1.00
0
2
4
6
8
10
12
Week

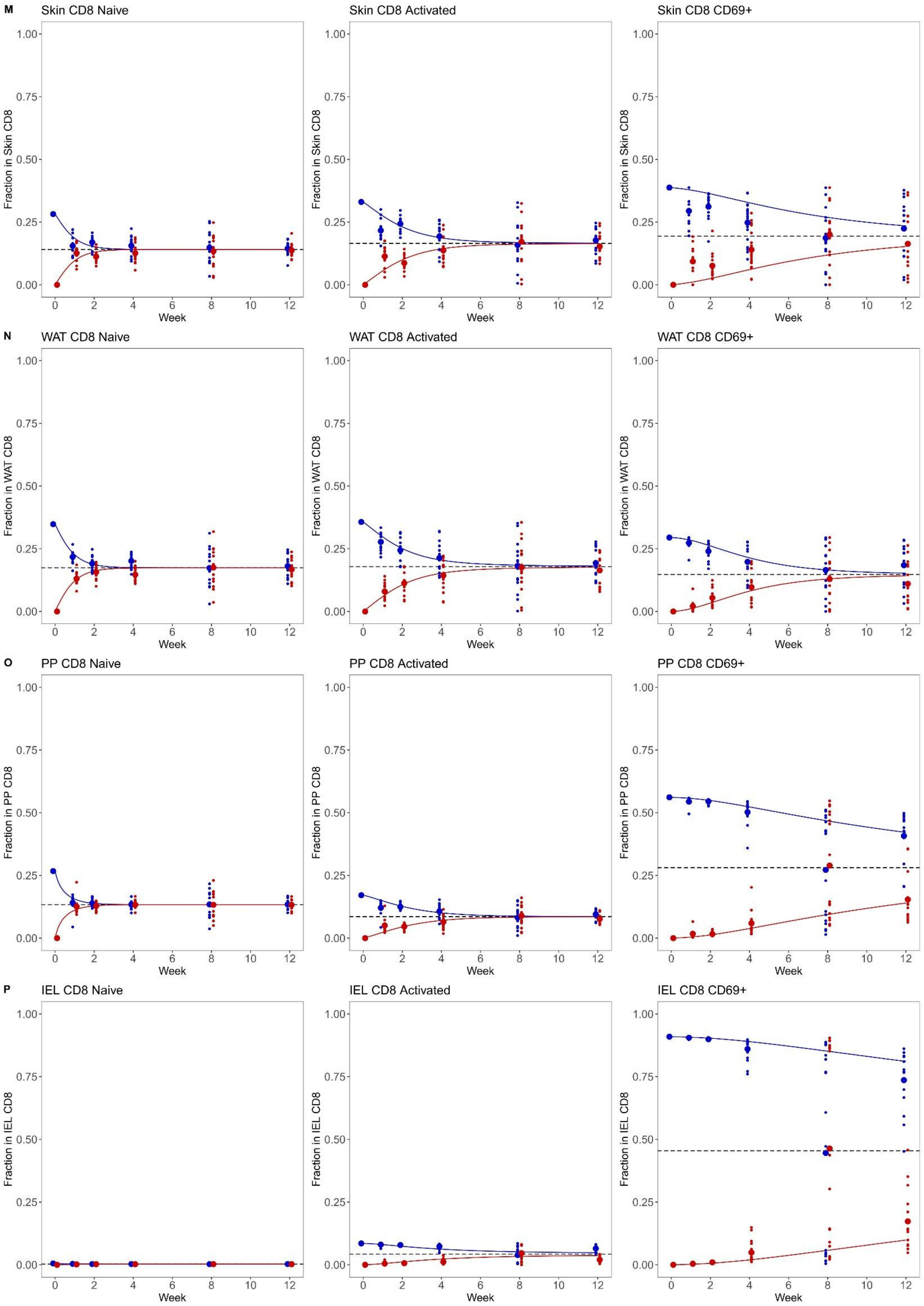

M
Skin CD8 Naive
Skin CD8 Activated
Skin CD8 CD69+
Fraction in Skin CD8
N
WAT CD8 Naive
WAT CD8 Activated
WAT CD8 CD69+
Fraction in WAT CD8
O
PP CD8 Naive
PP CD8 Activated
PP CD8 CD69+
Fraction in PP CD8
P
IEL CD8 Naive
IEL CD8 Activated
IEL CD8 CD69+
Fraction in IEL CD8
0.00
0.25
0.50
0.75
1.00
0
2
4
6
8
10
12
Week

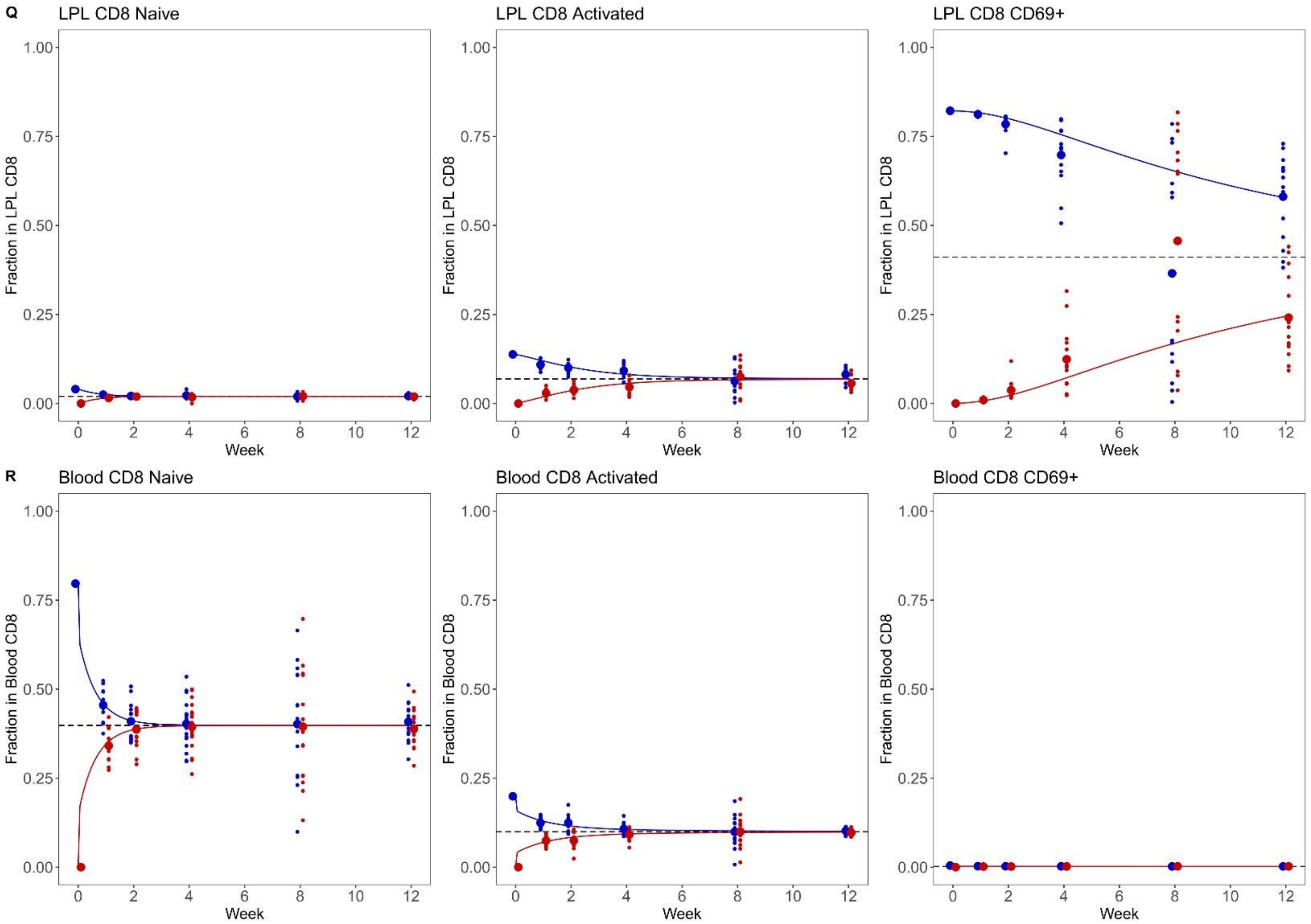

Q
LPL CD8 Naive
LPL CD8 Activated
LPL CD8 CD69+
Fraction in LPL CD8
R
Blood CD8 Naive
Blood CD8 Activated
Blood CD8 CD69+
Fraction in Blood CD8
1.00
0.75
0.50
0.25
0.00
Week
0 2 4 6 8 10 12

**Supplementary Figure 2. Markov chain models for tissue CD4 cellular kinetics.** CD45.1 mice were parabiosed to CD45.2 mice. Pairs of parabiotic animals were sacrificed at weeks 1, 2, 4, 8, and 12 for tissue analysis by flow cytometry (n=11,12,18,16,14). Markov chains were built to model the changes in cell state and tissue exchange, with each tissue built using a model containing the tissue, blood, and combined other tissues. Displayed are the original data points superimposed on the model predictions for naïve CD4 Tconv (left), activated CD4 Tconv (middle) and $CD69^+$ CD4 Tconv (right) from **A.** bone-marrow, **B.** spleen, **C.** LN, **D.** mLN, **E.** adrenals, **F.** brain, **G.** FRT, **H.** kidney, **I.** liver, **J.** lung, **K.** muscle, **L.** pancreas, **M.** skin, **N.** WAT, **O.** PP, **P.** IEL, **Q.** LPL, and **R.** blood. The blood model results displayed is from the spleen, blood and other tissues model, while all other tissues are displayed from their own model.

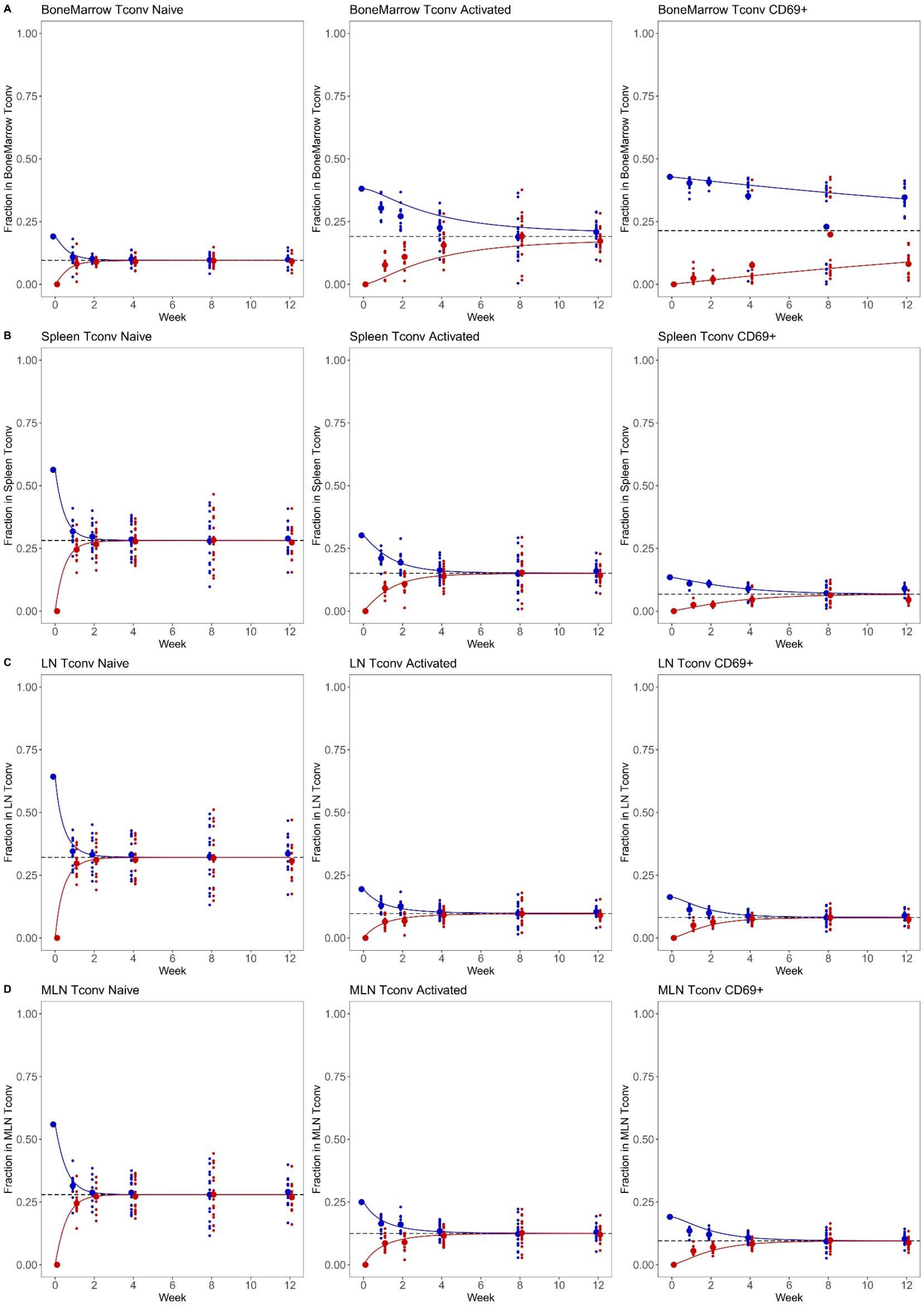
A
BoneMarrow Tconv Naive
BoneMarrow Tconv Activated
BoneMarrow Tconv CD69+
Fraction in BoneMarrow Tconv
B
Spleen Tconv Naive
Spleen Tconv Activated
Spleen Tconv CD69+
Fraction in Spleen Tconv
C
LN Tconv Naive
LN Tconv Activated
LN Tconv CD69+
Fraction in LN Tconv
D
MLN Tconv Naive
MLN Tconv Activated
MLN Tconv CD69+
Fraction in MLN Tconv
0.00
0.25
0.50
0.75
1.00
0
2
4
6
8
10
12
Week

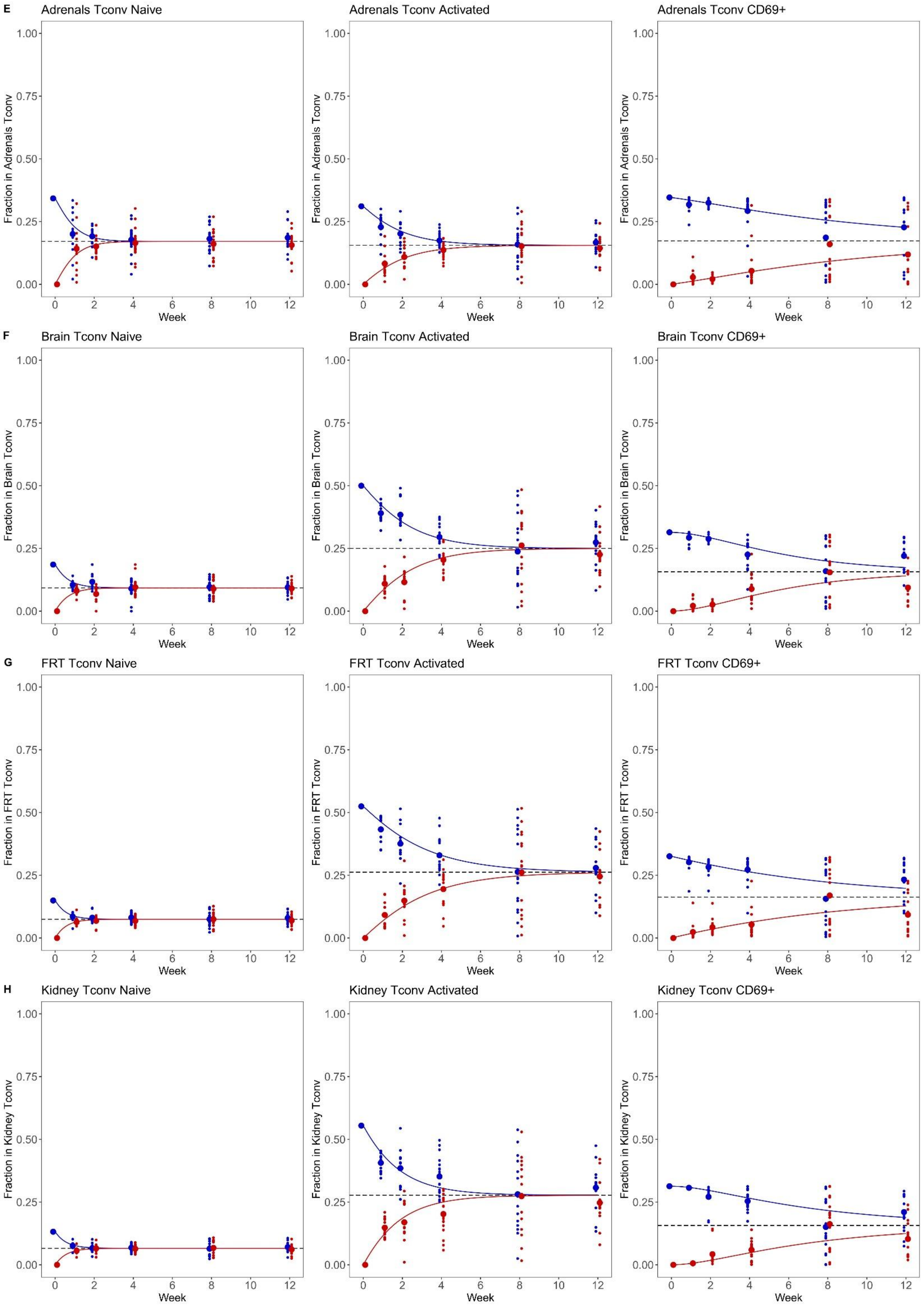
E
Adrenals Tconv Naive
Adrenals Tconv Activated
Adrenals Tconv CD69+
Fraction in Adrenals Tconv
F
Brain Tconv Naive
Brain Tconv Activated
Brain Tconv CD69+
Fraction in Brain Tconv
G
FRT Tconv Naive
FRT Tconv Activated
FRT Tconv CD69+
Fraction in FRT Tconv
H
Kidney Tconv Naive
Kidney Tconv Activated
Kidney Tconv CD69+
Fraction in Kidney Tconv
1.00
0.75
0.50
0.25
0.00
0
2
4
6
8
10
12
Week

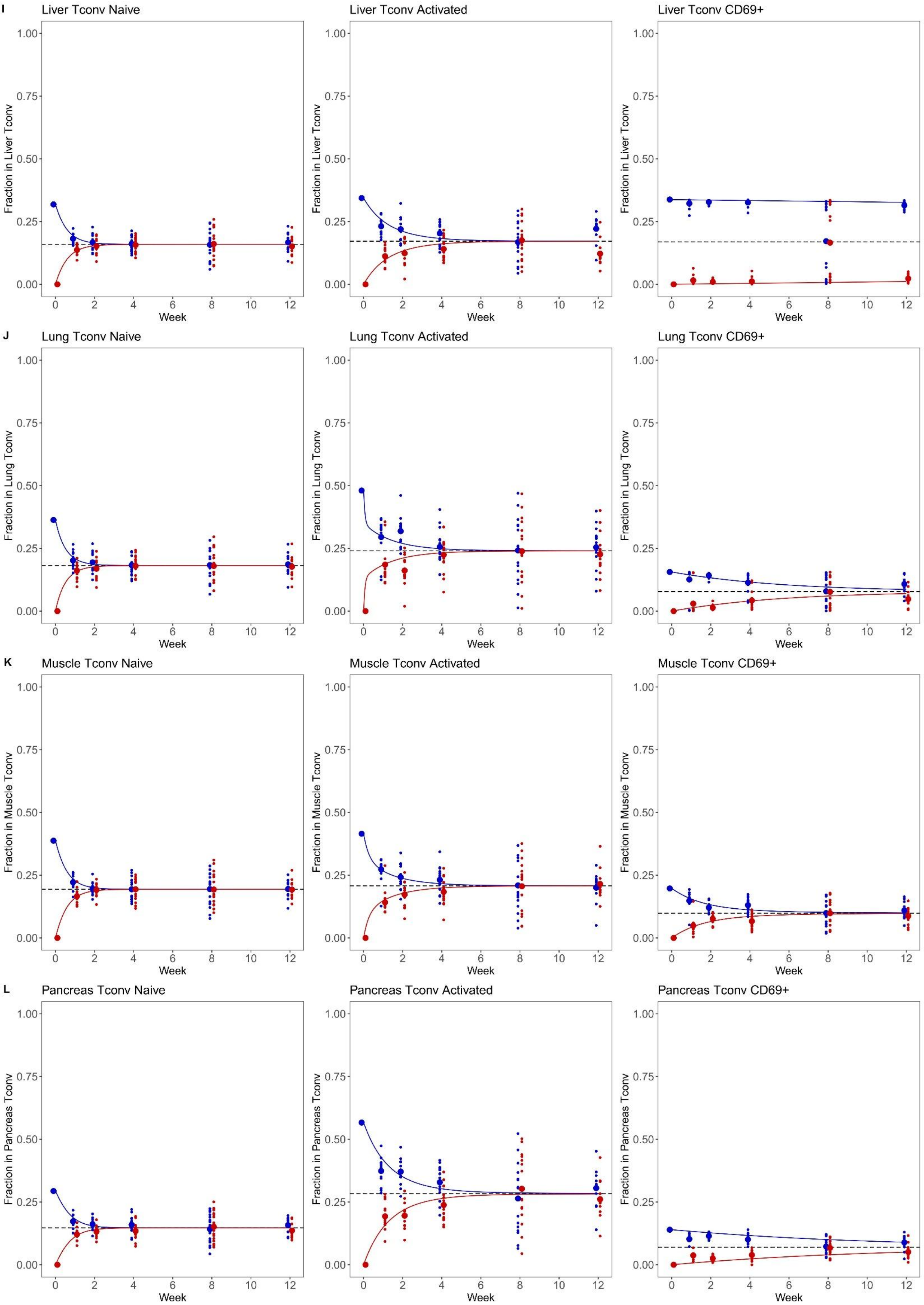

I
Liver Tconv Naive
Liver Tconv Activated
Liver Tconv CD69+
Fraction in Liver Tconv
J
Lung Tconv Naive
Lung Tconv Activated
Lung Tconv CD69+
Fraction in Lung Tconv
K
Muscle Tconv Naive
Muscle Tconv Activated
Muscle Tconv CD69+
Fraction in Muscle Tconv
L
Pancreas Tconv Naive
Pancreas Tconv Activated
Pancreas Tconv CD69+
Fraction in Pancreas Tconv
0.00
0.25
0.50
0.75
1.00
0
2
4
6
8
10
12
Week

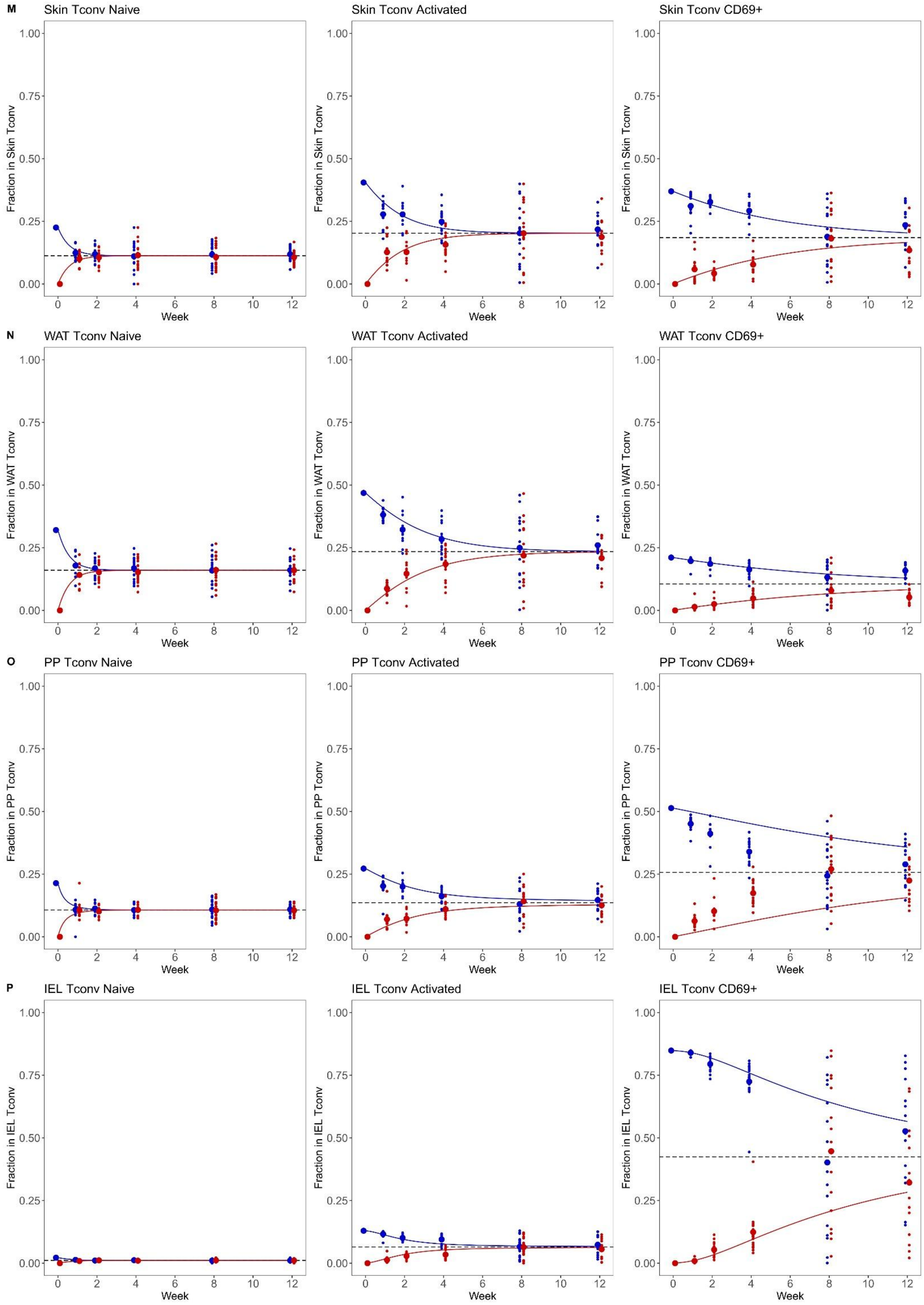
M
Skin Tconv Naive
Skin Tconv Activated
Skin Tconv CD69+
Fraction in Skin Tconv
N
WAT Tconv Naive
WAT Tconv Activated
WAT Tconv CD69+
Fraction in WAT Tconv
O
PP Tconv Naive
PP Tconv Activated
PP Tconv CD69+
Fraction in PP Tconv
P
IEL Tconv Naive
IEL Tconv Activated
IEL Tconv CD69+
Fraction in IEL Tconv
0.00
0.25
0.50
0.75
1.00
0
2
4
6
8
10
12
Week

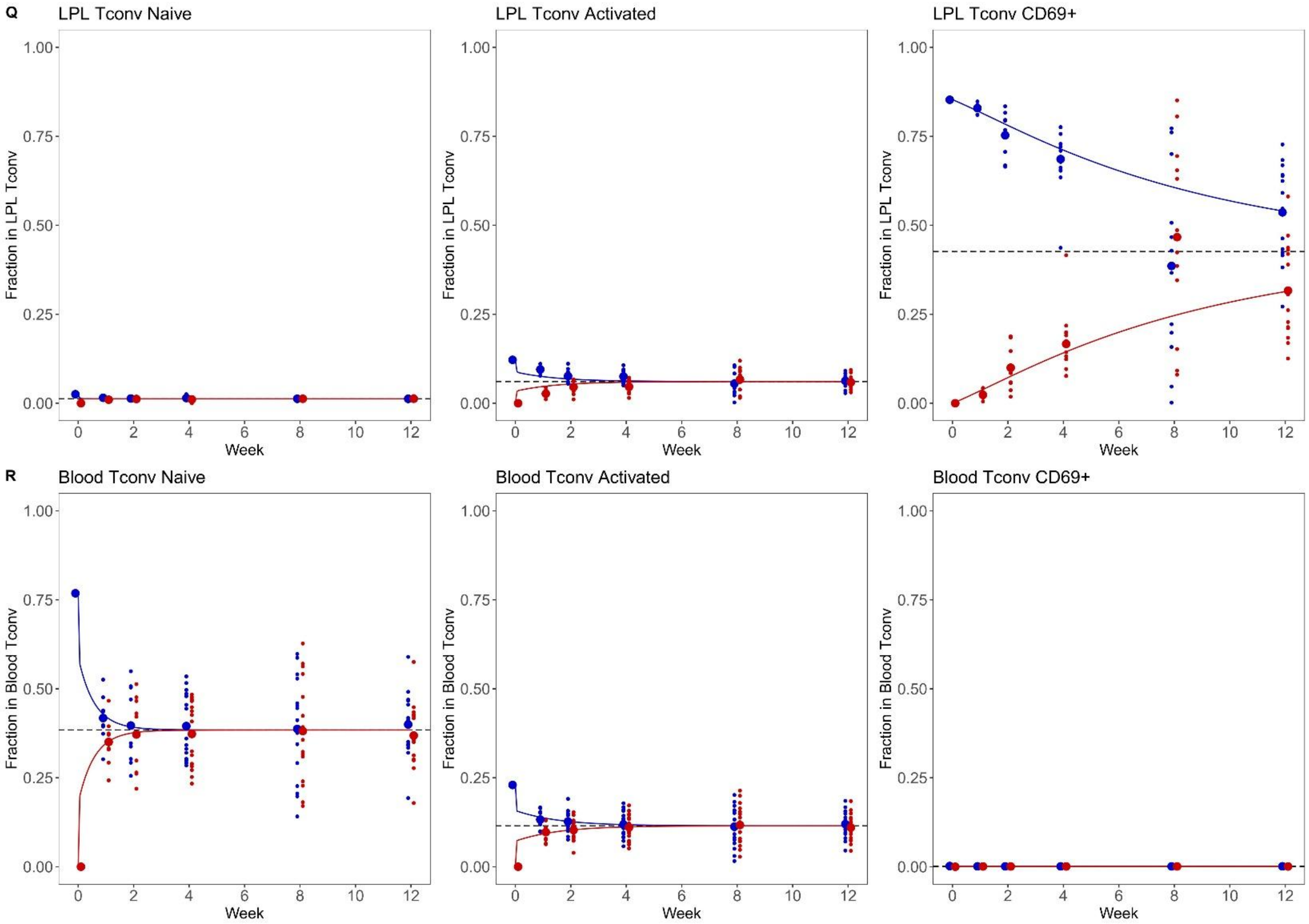

Q
LPL Tconv Naive
LPL Tconv Activated
LPL Tconv CD69+
Fraction in LPL Tconv
1.00
0.75
0.50
0.25
0.00
Week
0
2
4
6
8
10
12
R
Blood Tconv Naive
Blood Tconv Activated
Blood Tconv CD69+
Fraction in Blood Tconv

**Supplementary Figure 3. Markov chain models for tissue Treg cellular kinetics.** CD45.1 mice were parabiosed to CD45.2 mice. Pairs of parabiotic animals were sacrificed at weeks 1, 2, 4, 8, and 12 for tissue analysis by flow cytometry (n=11,12,18,16,14). Markov chains were built to model the changes in cell state and tissue exchange, with each tissue built using a model containing the tissue, blood, and combined other tissues. Displayed are the original data points superimposed on the model predictions for resting Treg (left), antigen-experienced Treg (middle) and $CD69^{+}$ Treg from **A.** bone-marrow, **B.** spleen, **C.** LN, **D.** mLN, **E.** adrenals, **F.** brain, **G.** FRT, **H.** kidney, **I.** liver, **J.** lung, **K.** muscle, **L.** pancreas, **M.** skin, **N.** WAT, **O.** PP, **P.** IEL, **Q.** LPL, and **R.** blood. The blood model results displayed is from the spleen, blood and other tissues model, while all other tissues are displayed from their own model.

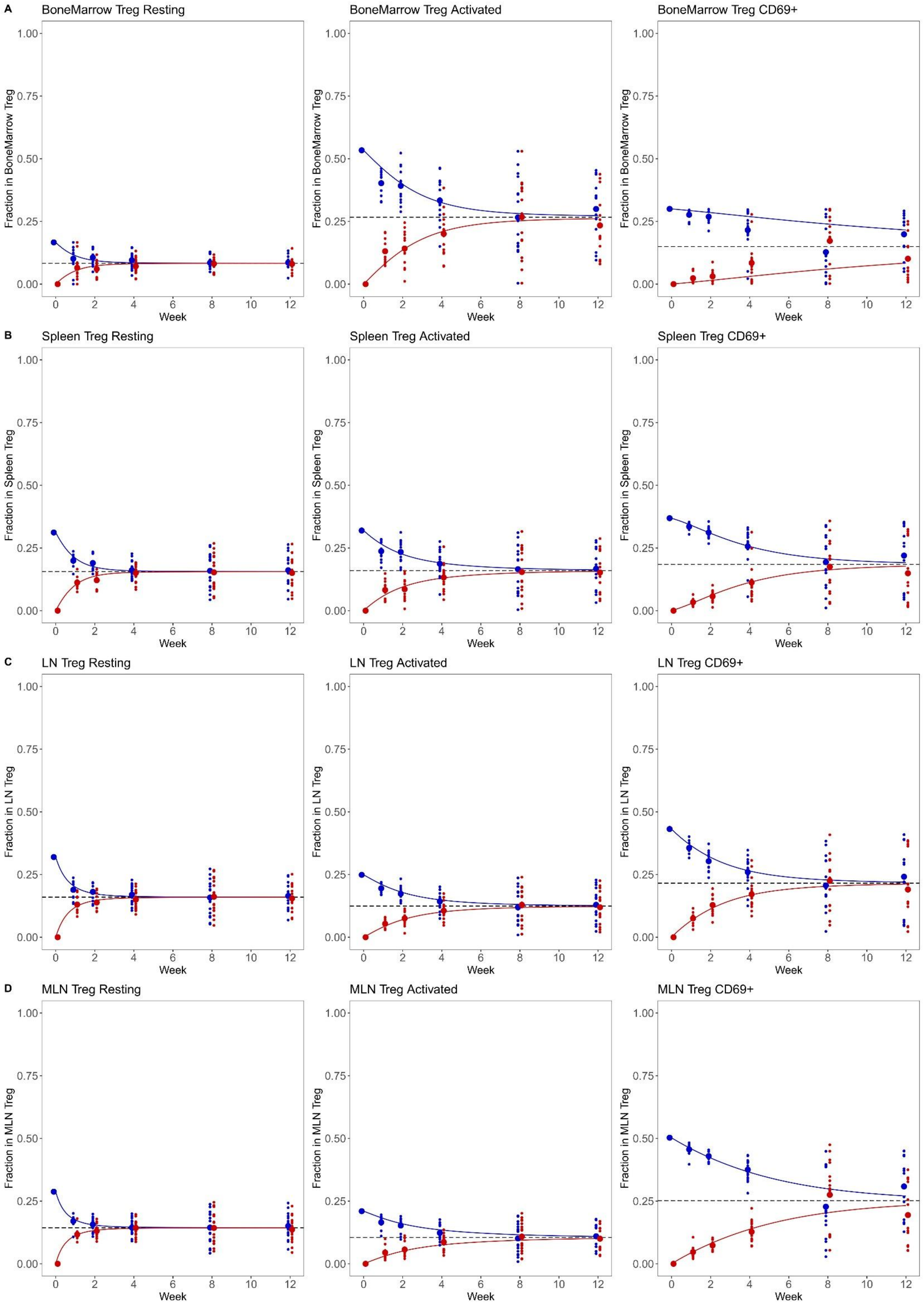

A
BoneMarrow Treg Resting
BoneMarrow Treg Activated
BoneMarrow Treg CD69+
Fraction in BoneMarrow Treg
B
Spleen Treg Resting
Spleen Treg Activated
Spleen Treg CD69+
Fraction in Spleen Treg
C
LN Treg Resting
LN Treg Activated
LN Treg CD69+
Fraction in LN Treg
D
MLN Treg Resting
MLN Treg Activated
MLN Treg CD69+
Fraction in MLN Treg
Week

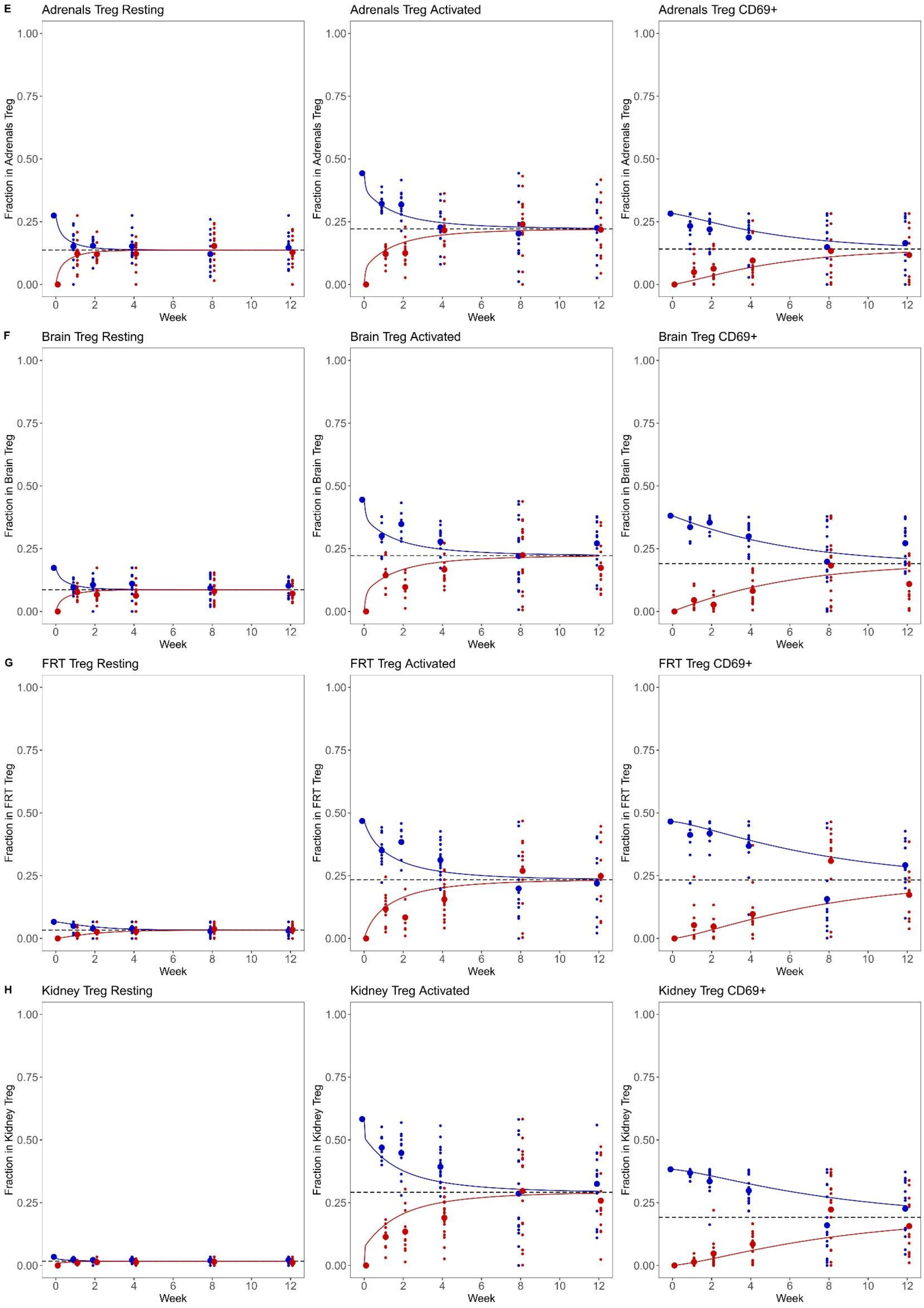

E
Adrenals Treg Resting
Adrenals Treg Activated
Adrenals Treg CD69+
Fraction in Adrenals Treg
F
Brain Treg Resting
Brain Treg Activated
Brain Treg CD69+
Fraction in Brain Treg
G
FRT Treg Resting
FRT Treg Activated
FRT Treg CD69+
Fraction in FRT Treg
H
Kidney Treg Resting
Kidney Treg Activated
Kidney Treg CD69+
Fraction in Kidney Treg
1.00
0.75
0.50
0.25
0.00
0
2
4
6
8
10
12
Week

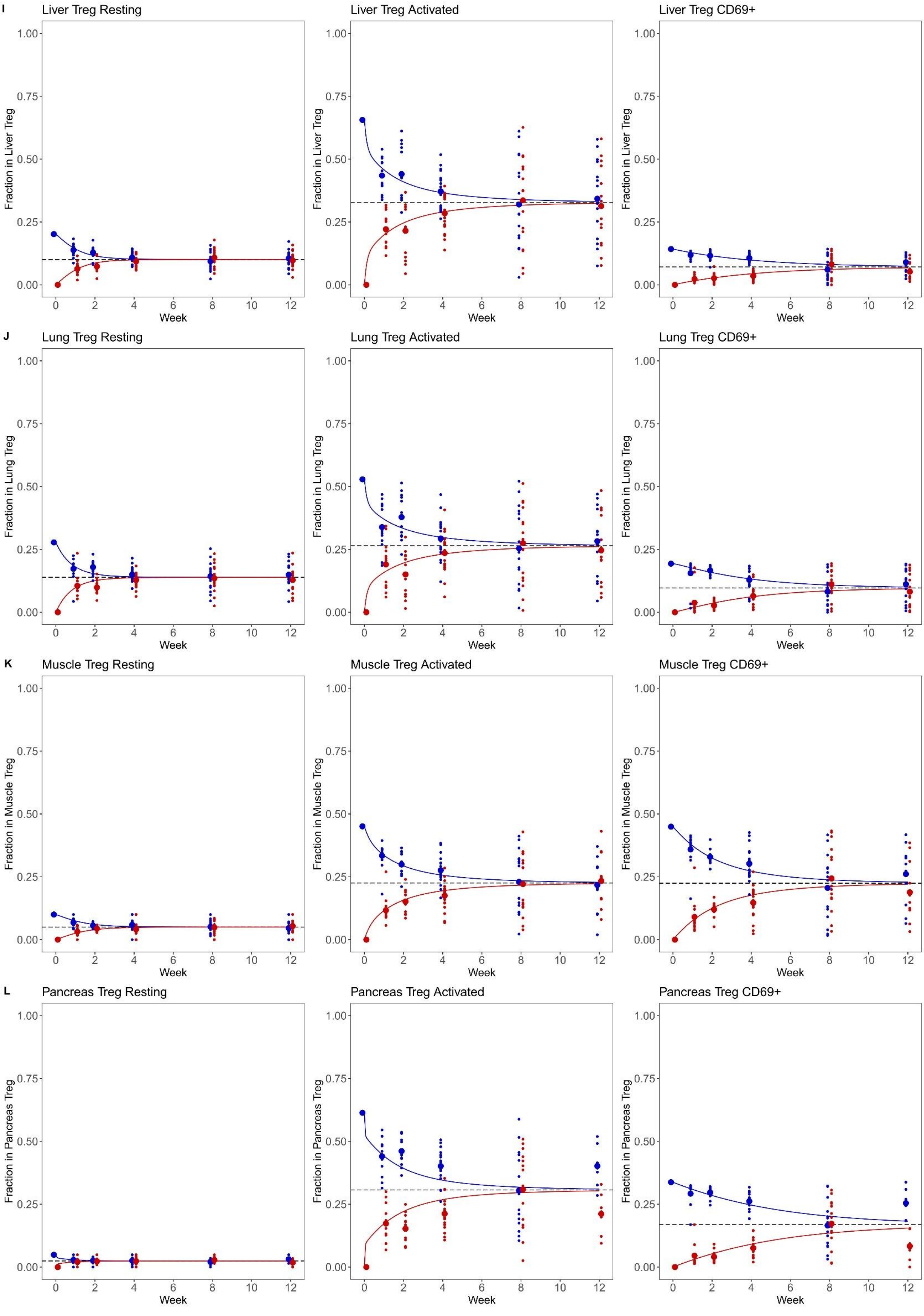

I
Liver Treg Resting
Liver Treg Activated
Liver Treg CD69+
Fraction in Liver Treg
J
Lung Treg Resting
Lung Treg Activated
Lung Treg CD69+
Fraction in Lung Treg
K
Muscle Treg Resting
Muscle Treg Activated
Muscle Treg CD69+
Fraction in Muscle Treg
L
Pancreas Treg Resting
Pancreas Treg Activated
Pancreas Treg CD69+
Fraction in Pancreas Treg
0.00
0.25
0.50
0.75
1.00
0
2
4
6
8
10
12
Week

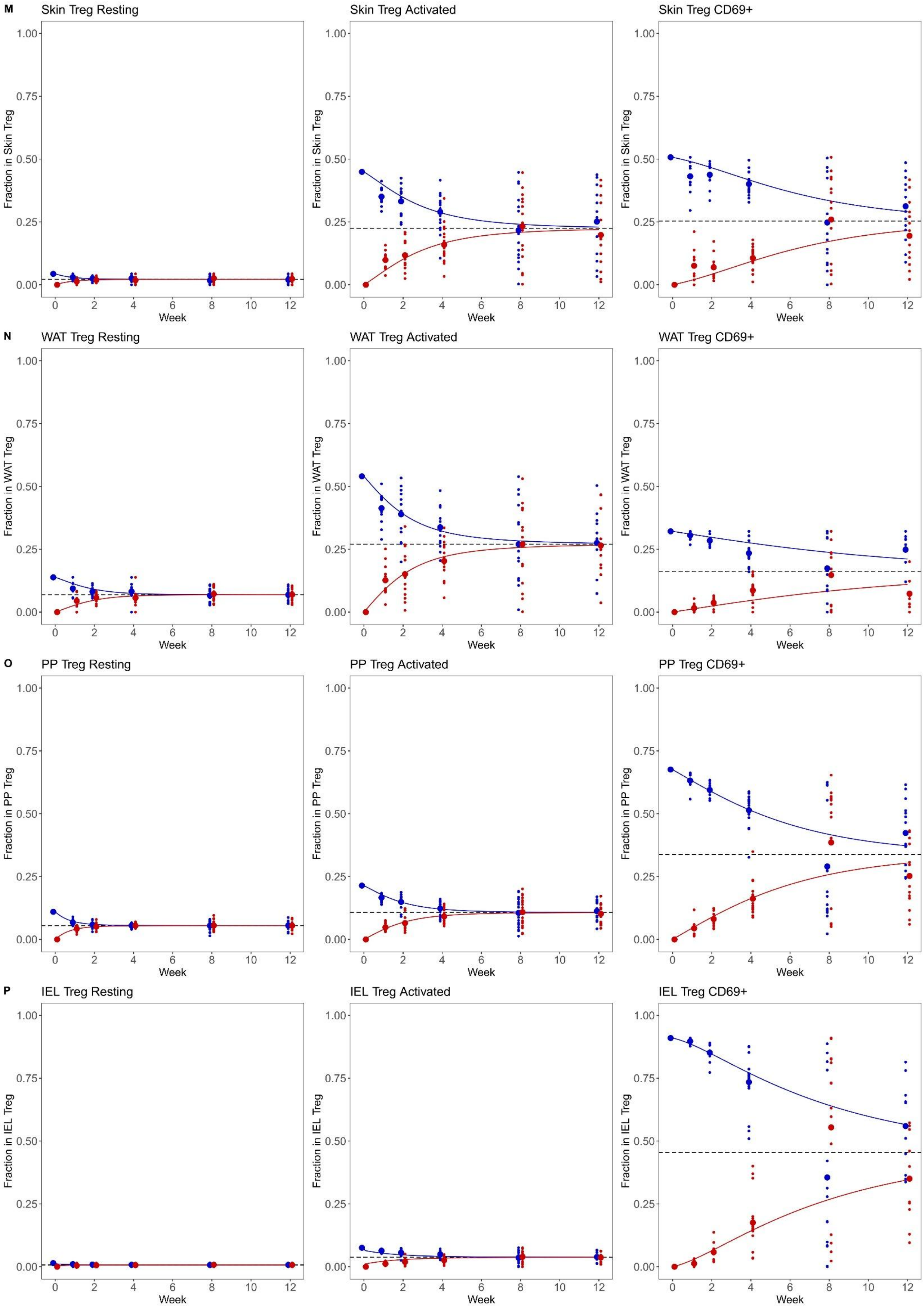
M
Skin Treg Resting
Skin Treg Activated
Skin Treg CD69+
Fraction in Skin Treg
N
WAT Treg Resting
WAT Treg Activated
WAT Treg CD69+
Fraction in WAT Treg
O
PP Treg Resting
PP Treg Activated
PP Treg CD69+
Fraction in PP Treg
P
IEL Treg Resting
IEL Treg Activated
IEL Treg CD69+
Fraction in IEL Treg
Week
0.00
0.25
0.50
0.75
1.00
0
2
4
6
8
10
12

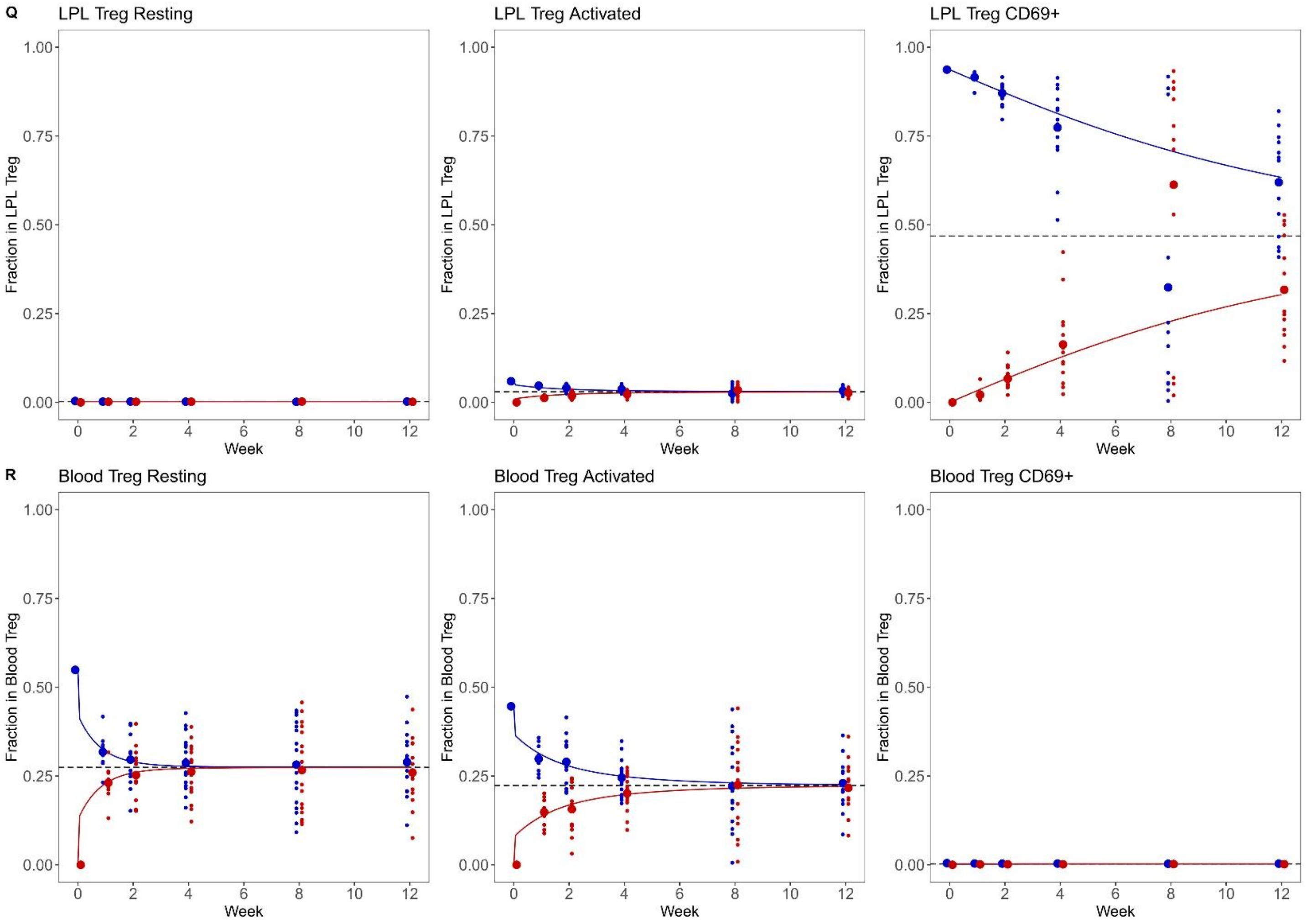

Q
LPL Treg Resting
LPL Treg Activated
LPL Treg CD69+
Fraction in LPL Treg
1.00
0.75
0.50
0.25
0.00
0
2
4
6
8
10
12
Week
R
Blood Treg Resting
Blood Treg Activated
Blood Treg CD69+
Fraction in Blood Treg

**Supplementary Figure 4. Numerical contribution to T cell populations in the tissue.** Source of cumulative entry per tissue, based on activation status of blood-based precursor, across bone marrow, lymphoid tissues (spleen, LN, MLN), non-lymphoid tissues (adrenals, brain, kidney, FRT, lung, liver, muscle, WAT, skin, pancreas) and gut-associated tissues (LPL, IEL, PP) and across cell states (naïve/resting, activated/antigen-experienced, $CD69^{+}$). Modelling data from **A.** CD8, **B.** CD4 Tconv, and **C.** CD4 Treg cells.

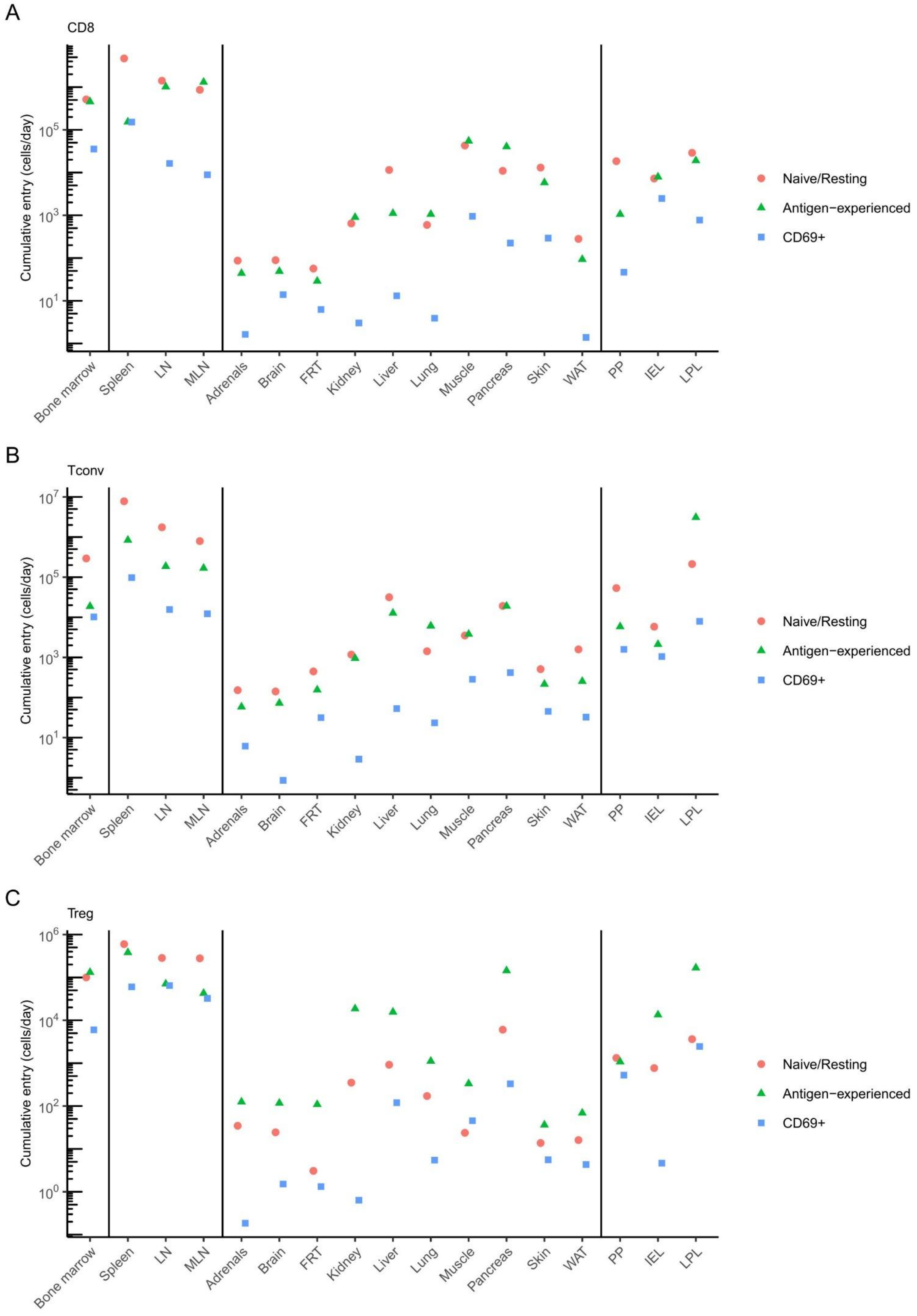

A
CD8
B
Tconv
C
Treg
Cumulative entry (cells/day)
$10^1$
$10^3$
$10^5$
$10^7$
$10^0$
$10^2$
$10^4$
$10^6$
Bone marrow
Spleen
LN
MLN
Adrenals
Brain
FRT
Kidney
Liver
Lung
Muscle
Pancreas
Skin
WAT
PP
IEL
LPL
Naive/Resting
Antigen−experienced
CD69+

**Supplementary Figure 5. Cellular longevity of tissue-trafficking T cells.** Synthetic T cells, as an in silico population obeying the probabilistic rules generated from the best-fit model of the empirical data, were simulated until cell "death". 10,000 simulated CD69$^{+}$ cells, distributed among tissues based on steady-state proportions, were simulated through to cell death, with a histogram of the distribution of durations of cellular survival. Histograms represent survival time of **(A)** CD8 **(B)** CD4 Tconv and **(C)** CD4 Treg cells, with highlighted median survival time. Cellular survival was cropped at 1000 days to represent cells reaching the typical lifespan of the animal.

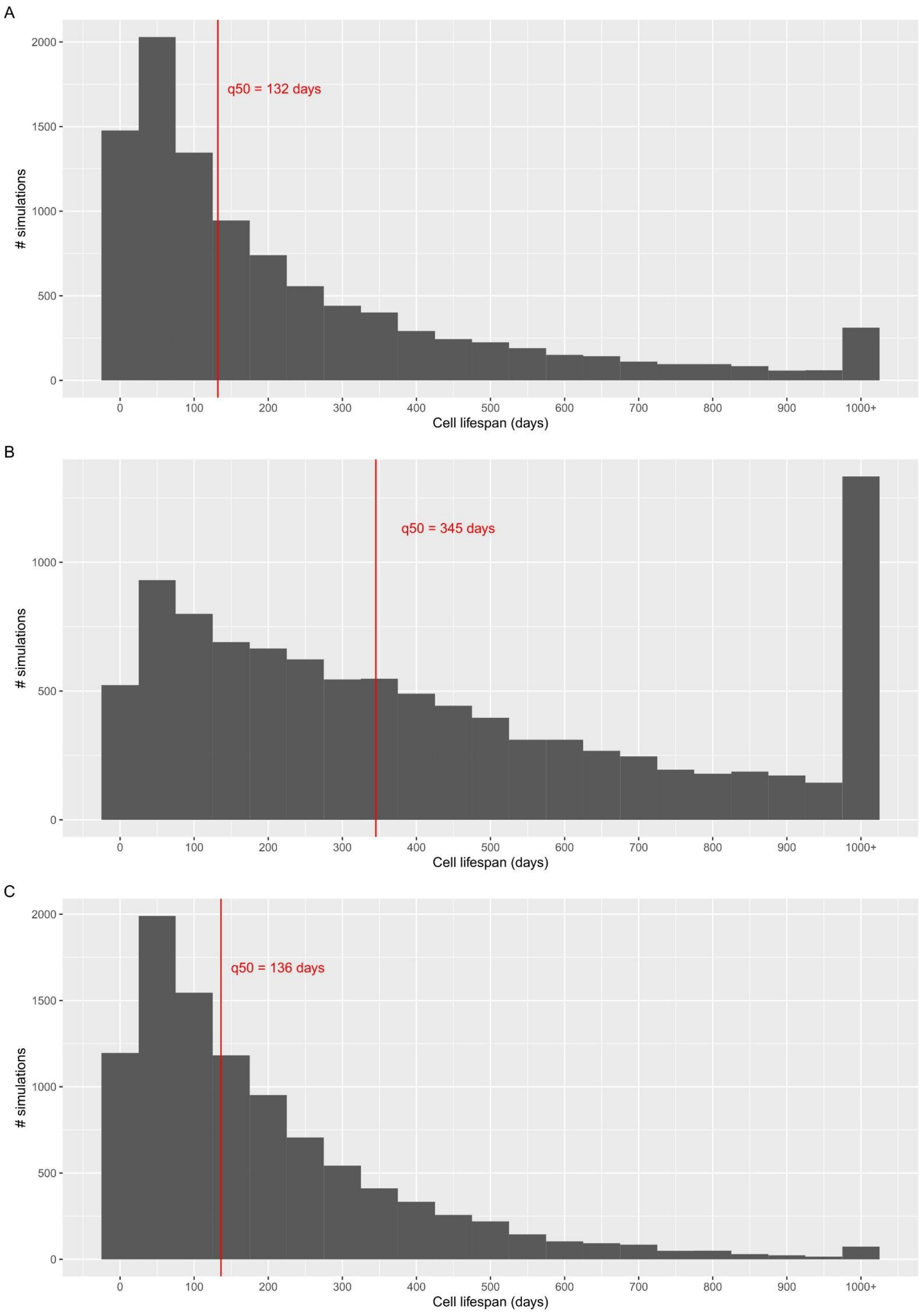

A
q50 = 132 days
### simulations
Cell lifespan (days)
B
q50 = 345 days
### simulations
Cell lifespan (days)
C
q50 = 136 days
### simulations
Cell lifespan (days)

**Supplementary Figure 6. Markov chain models for tissue B cell cellular kinetics.** CD45.1 mice were parabiosed to CD45.2 mice. Pairs of parabiotic animals were sacrificed at weeks 1, 2, 4, 8, and 12 for tissue analysis by flow cytometry (n=11,12,18,16,14). Markov chains were built to model the changes in cell state and tissue exchange, with each tissue built using a model containing the tissue, blood, and combined other tissues. Displayed are the original data points superimposed on the model predictions for naïve B cells (left), activated B cells (middle) and $CD69^{+}$ B cells (right) from **A.** spleen, **B.** LN, **C.** mLN, **D.** adrenals, **E.** brain, **F.** FRT, **G.** kidney, **H.** liver, **I.** lung, **J.** muscle, **K.** pancreas, **L.** skin, **M.** WAT, **N.** PP, **O.** IEL, **P.** LPL, and **Q.** blood. Bone-marrow was excluded from analysis, as the site for primary B cell differentiation. The blood model results displayed is from the spleen, blood and other tissues model, while all other tissues are displayed from their own model.

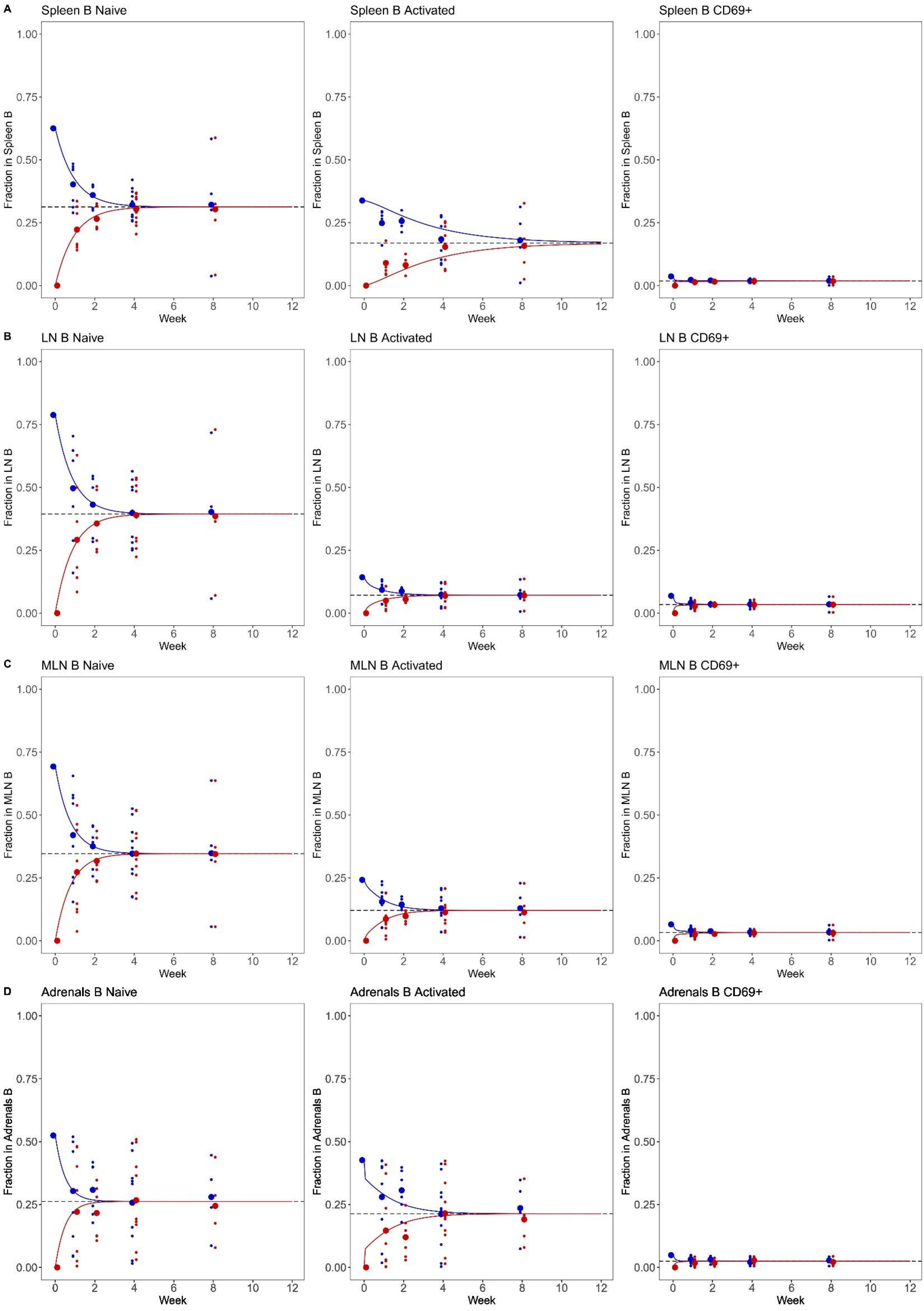

A
Spleen B Naive
Spleen B Activated
Spleen B CD69+
Fraction in Spleen B
B
LN B Naive
LN B Activated
LN B CD69+
Fraction in LN B
C
MLN B Naive
MLN B Activated
MLN B CD69+
Fraction in MLN B
D
Adrenals B Naive
Adrenals B Activated
Adrenals B CD69+
Fraction in Adrenals B
Week
0.00
0.25
0.50
0.75
1.00
0
2
4
6
8
10
12

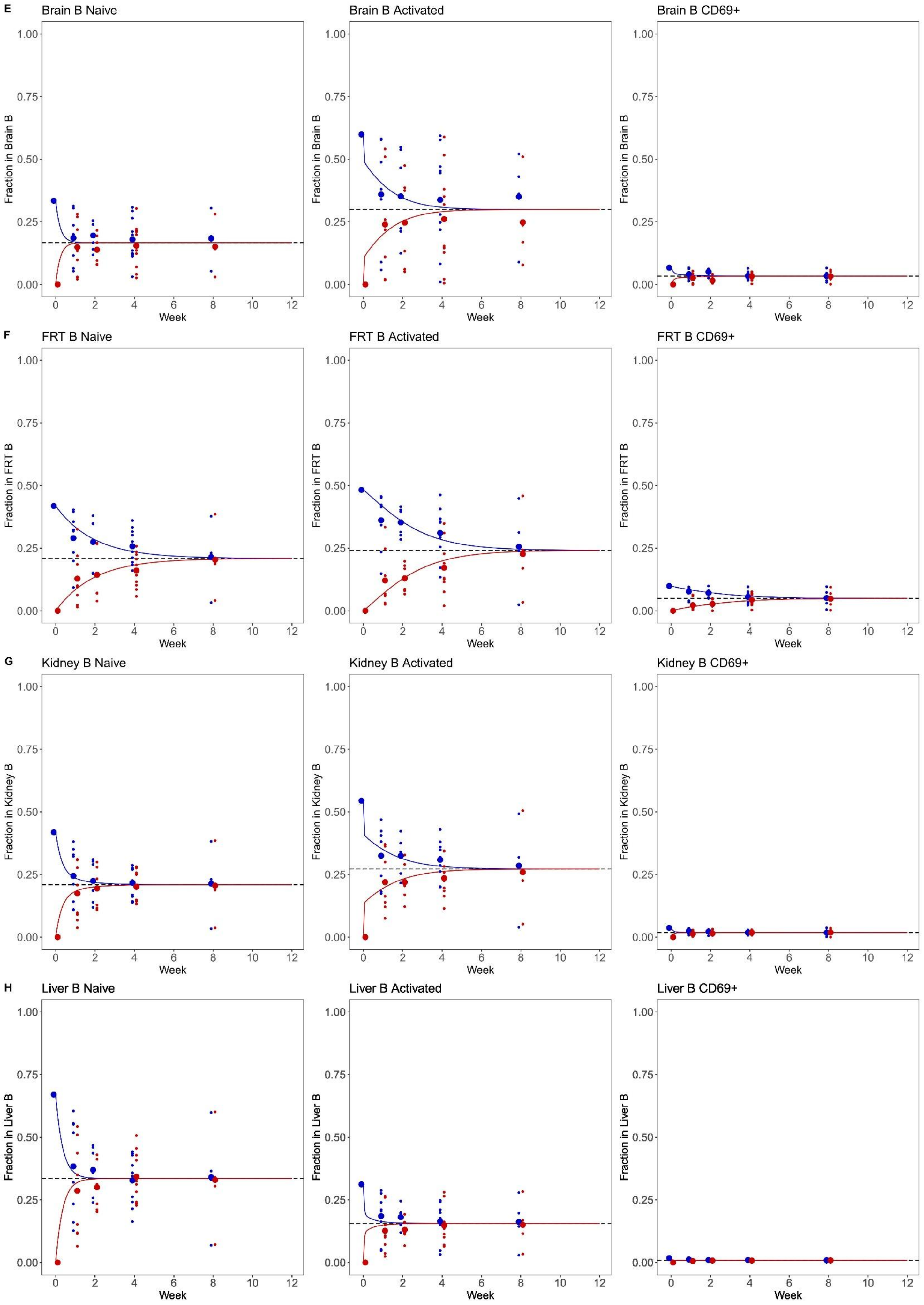

E
Brain B Naive
Brain B Activated
Brain B CD69+
Fraction in Brain B
F
FRT B Naive
FRT B Activated
FRT B CD69+
Fraction in FRT B
G
Kidney B Naive
Kidney B Activated
Kidney B CD69+
Fraction in Kidney B
H
Liver B Naive
Liver B Activated
Liver B CD69+
Fraction in Liver B
0.00
0.25
0.50
0.75
1.00
0
2
4
6
8
10
12
Week

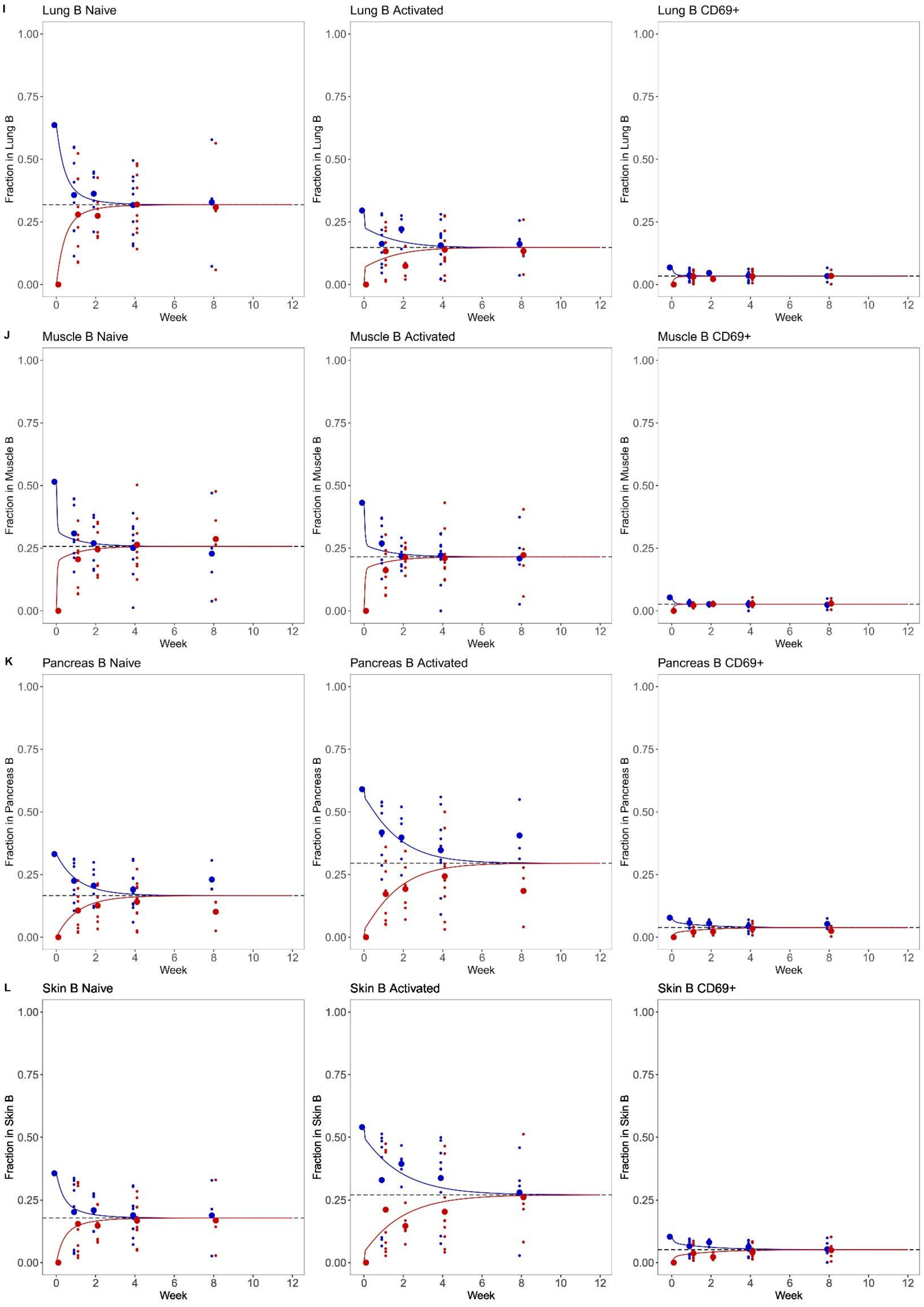

I
Lung B Naive
Lung B Activated
Lung B CD69+
Fraction in Lung B
J
Muscle B Naive
Muscle B Activated
Muscle B CD69+
Fraction in Muscle B
K
Pancreas B Naive
Pancreas B Activated
Pancreas B CD69+
Fraction in Pancreas B
L
Skin B Naive
Skin B Activated
Skin B CD69+
Fraction in Skin B
0.00
0.25
0.50
0.75
1.00
0
2
4
6
8
10
12
Week

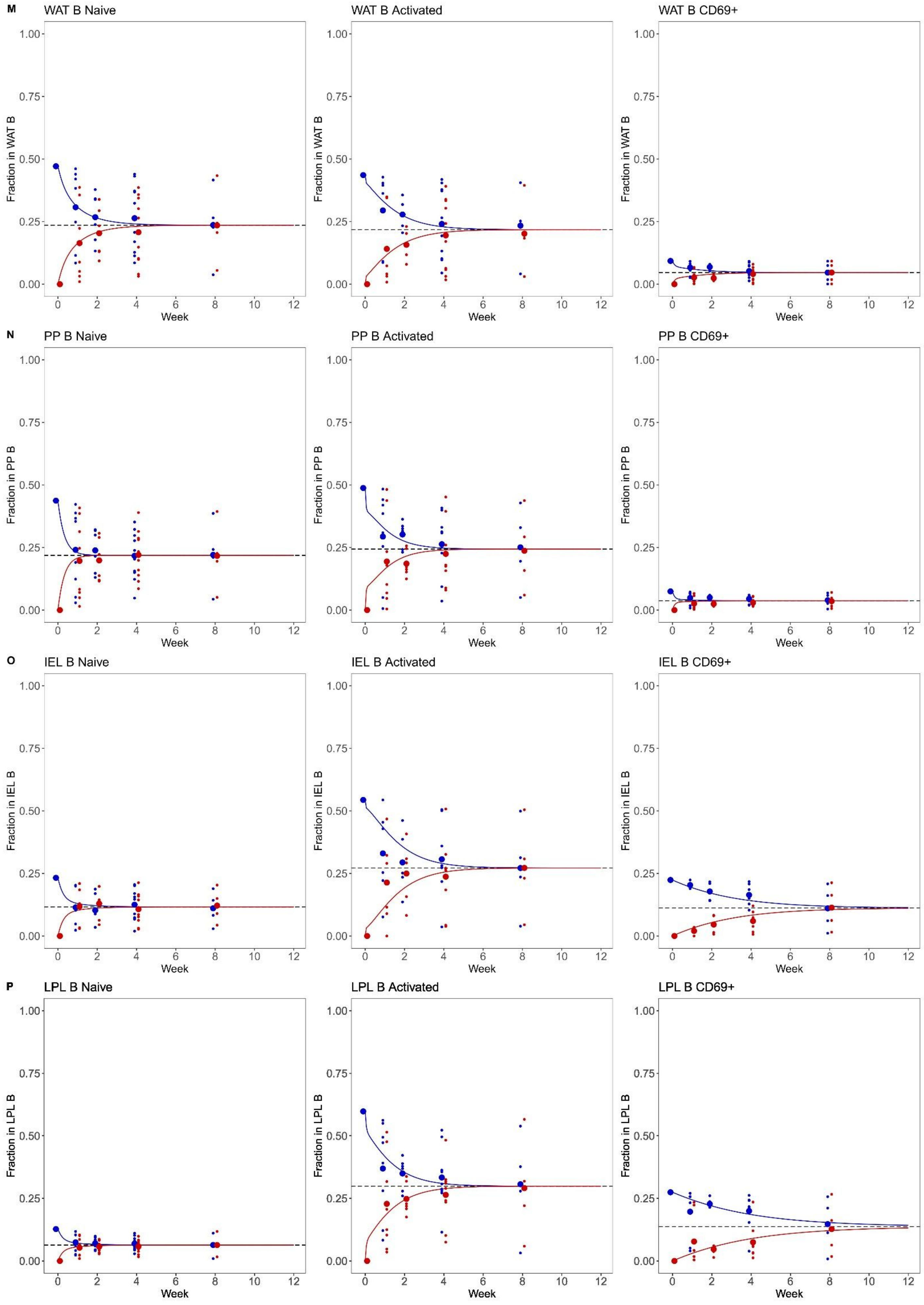

M
WAT B Naive
WAT B Activated
WAT B CD69+
Fraction in WAT B
N
PP B Naive
PP B Activated
PP B CD69+
Fraction in PP B
O
IEL B Naive
IEL B Activated
IEL B CD69+
Fraction in IEL B
P
LPL B Naive
LPL B Activated
LPL B CD69+
Fraction in LPL B
Week
0.00
0.25
0.50
0.75
1.00
0
2
4
6
8
10
12

**Q**

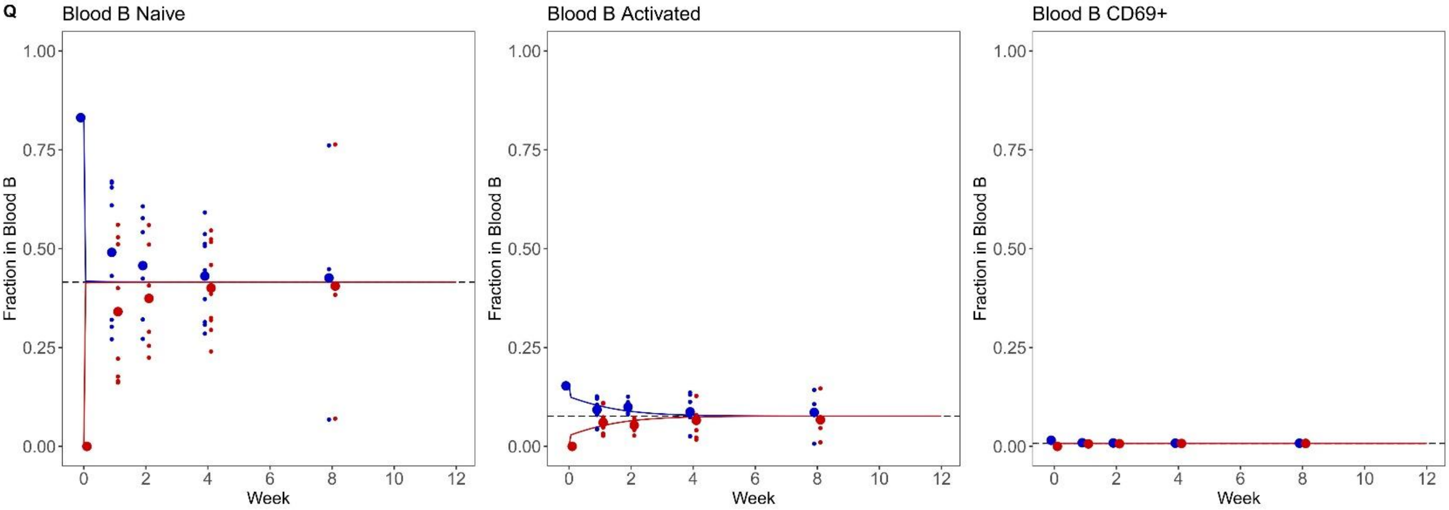

Blood B Naive
Blood B Activated
Blood B CD69+
Fraction in Blood B
Week
1.00
0.75
0.50
0.25
0.00
0
2
4
6
8
10
12

**Supplementary Figure 7. Markov chain models for tissue NK cell cellular kinetics.** CD45.1 mice were parabiosed to CD45.2 mice. Pairs of parabiotic animals were sacrificed at weeks 1, 2, 4, 8, and 12 for tissue analysis by flow cytometry (n=11,12,18,16,14). Markov chains were built to model the changes in cell state and tissue exchange, with each tissue built using a model containing the tissue, blood, and combined other tissues. Displayed are the original data points superimposed on the model predictions for naïve NK cells (left), activated NK cells (middle) and $CD69^+$ NK cells (right) from **A.** spleen, **B.** LN, **C.** mLN, **D.** adrenals, **E.** brain, **F.** FRT, **G.** kidney, **H.** liver, **I.** lung, **J.** muscle, **K.** pancreas, **L.** skin, **M.** WAT, **N.** PP, **O.** IEL, **P.** LPL, and **Q.** blood. Bone-marrow was excluded from analysis, as the site for primary NK cell differentiation. The blood model results displayed is from the spleen, blood and other tissues model, while all other tissues are displayed from their own model.

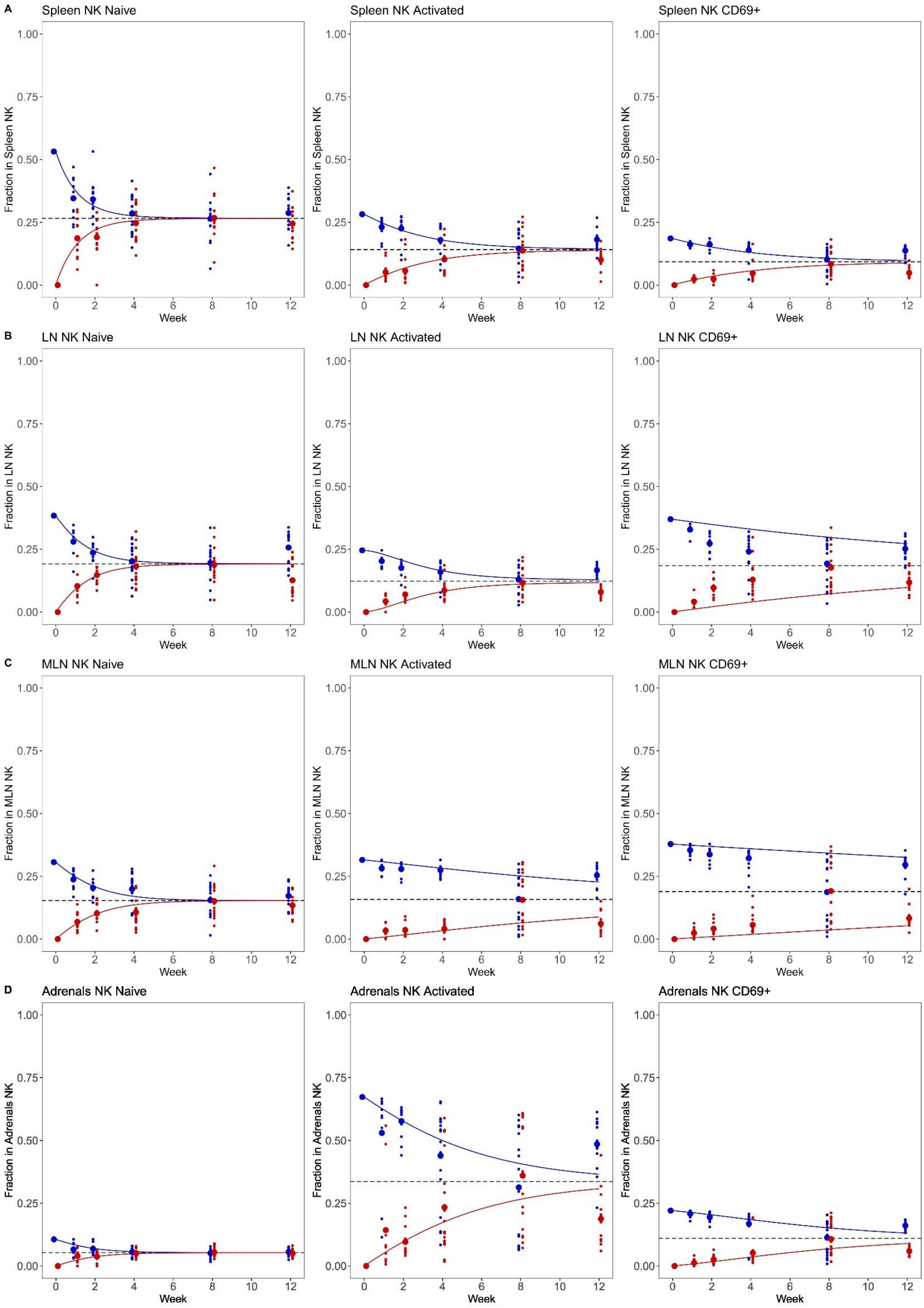
A
Spleen NK Naive
Spleen NK Activated
Spleen NK CD69+
Fraction in Spleen NK
B
LN NK Naive
LN NK Activated
LN NK CD69+
Fraction in LN NK
C
MLN NK Naive
MLN NK Activated
MLN NK CD69+
Fraction in MLN NK
D
Adrenals NK Naive
Adrenals NK Activated
Adrenals NK CD69+
Fraction in Adrenals NK
Week
0.00
0.25
0.50
0.75
1.00
0
2
4
6
8
10
12

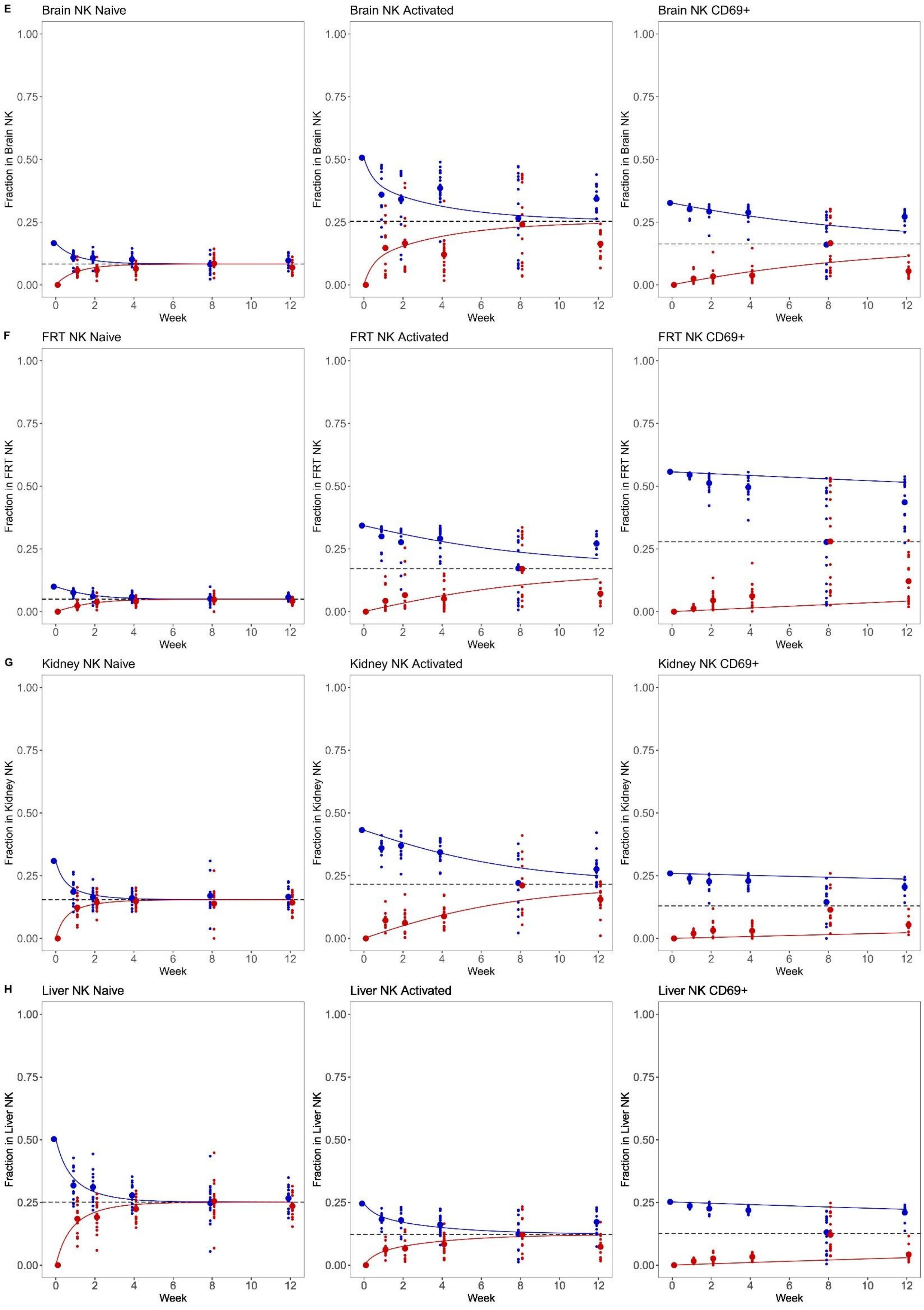
E
Brain NK Naive
Brain NK Activated
Brain NK CD69+
Fraction in Brain NK
F
FRT NK Naive
FRT NK Activated
FRT NK CD69+
Fraction in FRT NK
G
Kidney NK Naive
Kidney NK Activated
Kidney NK CD69+
Fraction in Kidney NK
H
Liver NK Naive
Liver NK Activated
Liver NK CD69+
Fraction in Liver NK
0.00
0.25
0.50
0.75
1.00
0
2
4
6
8
10
12
Week

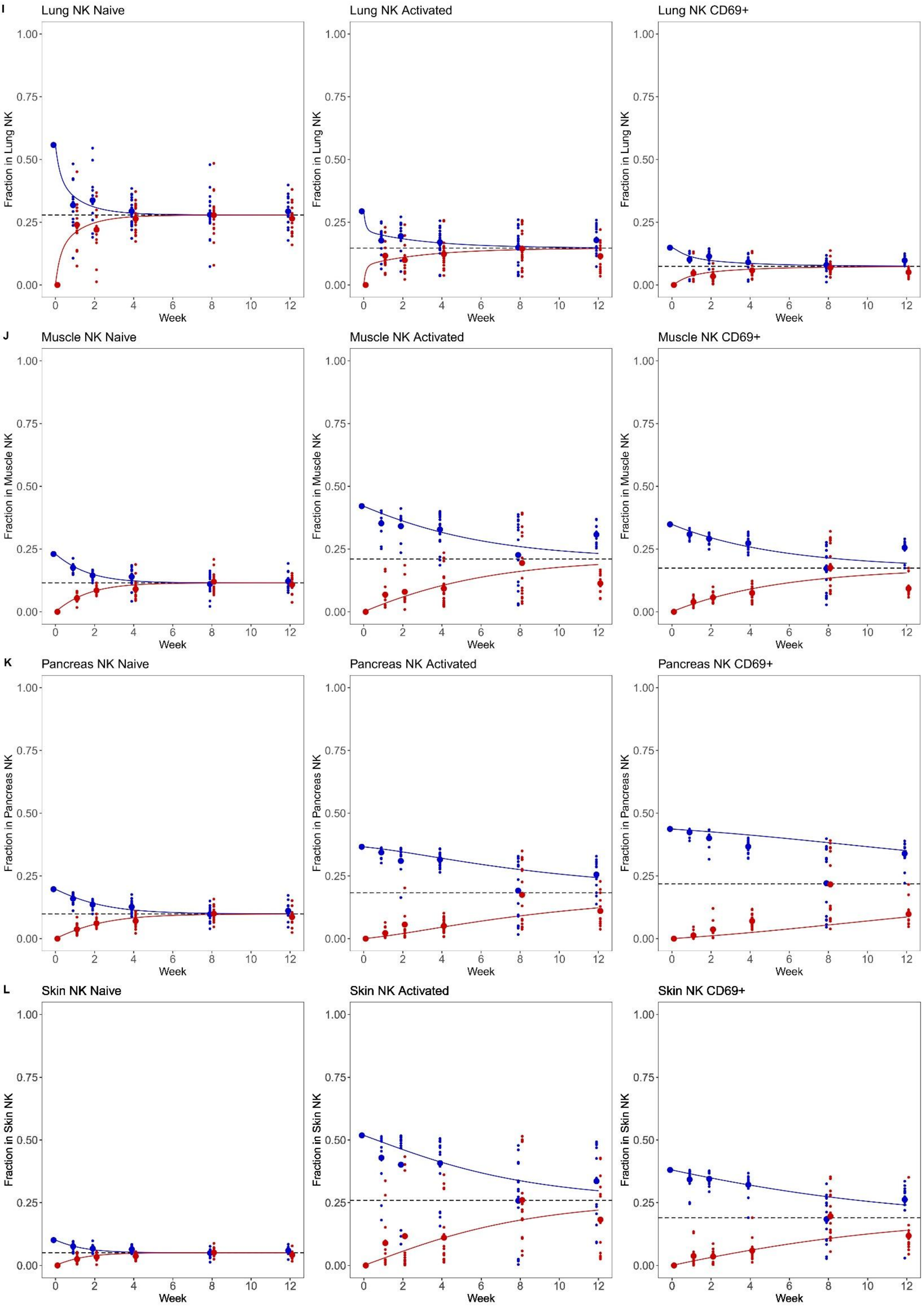

I
Lung NK Naive
Lung NK Activated
Lung NK CD69+
Fraction in Lung NK
J
Muscle NK Naive
Muscle NK Activated
Muscle NK CD69+
Fraction in Muscle NK
K
Pancreas NK Naive
Pancreas NK Activated
Pancreas NK CD69+
Fraction in Pancreas NK
L
Skin NK Naive
Skin NK Activated
Skin NK CD69+
Fraction in Skin NK
0.00
0.25
0.50
0.75
1.00
0
2
4
6
8
10
12
Week

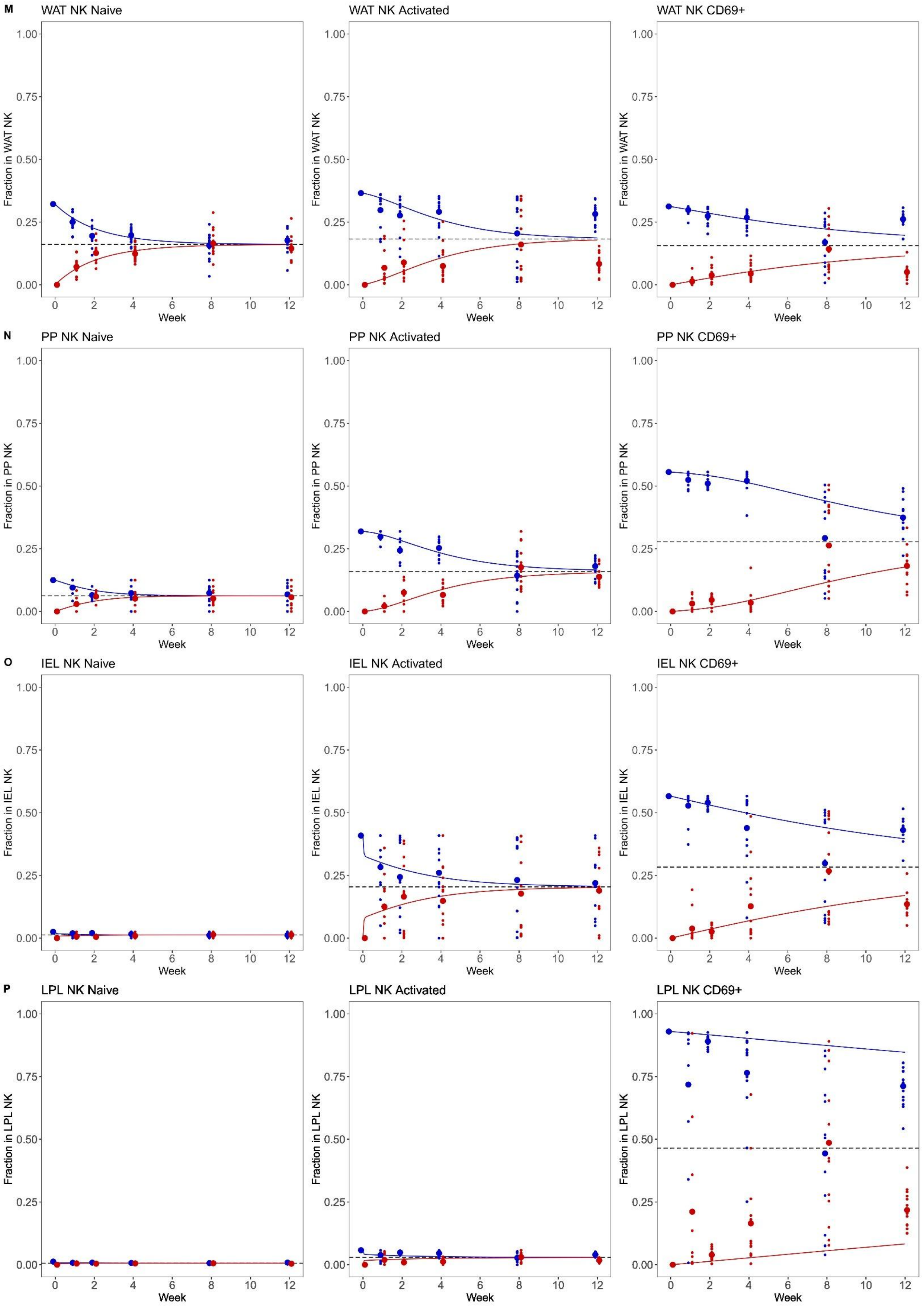
M
WAT NK Naive
WAT NK Activated
WAT NK CD69+
Fraction in WAT NK
N
PP NK Naive
PP NK Activated
PP NK CD69+
Fraction in PP NK
O
IEL NK Naive
IEL NK Activated
IEL NK CD69+
Fraction in IEL NK
P
LPL NK Naive
LPL NK Activated
LPL NK CD69+
Fraction in LPL NK
0.00
0.25
0.50
0.75
1.00
0
2
4
6
8
10
12
Week

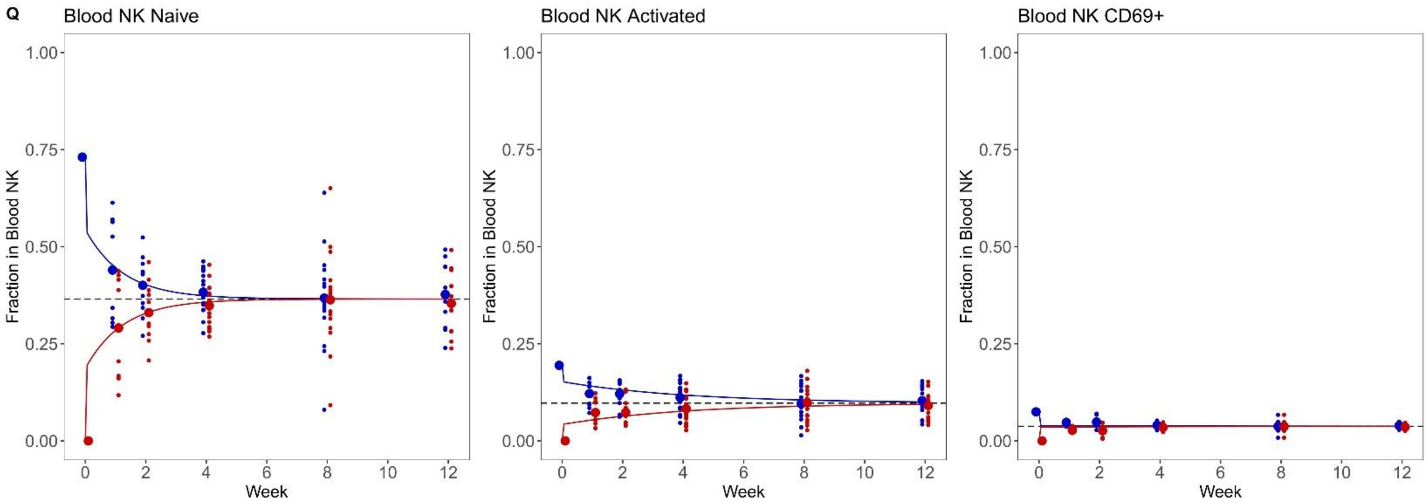
Q
Blood NK Naive
Blood NK Activated
Blood NK CD69+
Fraction in Blood NK
1.00
0.75
0.50
0.25
0.00
Week
0
2
4
6
8
10
12
